# Sensory axons induce epithelial lipid microdomain remodeling and determine the distribution of junctions in the epidermis

**DOI:** 10.1101/2021.12.23.474011

**Authors:** Jeffrey B. Rosa, Khaled Y. Nassman, Alvaro Sagasti

## Abstract

Epithelial cell properties are determined by the polarized distribution of membrane lipids, the cytoskeleton, and adhesive junctions. Epithelia are often profusely innervated, but little work has addressed how contact with neurites affects the polarized organization of epithelial components. In previous work, we found that basal keratinocytes in the larval zebrafish epidermis wrap around axons to enclose them in ensheathment channels sealed by autotypic cell junctions. In this study, we used live imaging to characterize how sensory axons remodel cell membranes, the actin cytoskeleton, and adhesive junctions in basal keratinocytes. At the apical surface of basal keratinocytes, axons promoted the formation of lipid microdomains quantitatively enriched in reporters for PI(4,5)P2 and liquid-ordered (Lo) membranes. Lipid microdomains supported the formation of cadherin-enriched F-actin protrusions, which wrapped around axons, likely initiating the formation of ensheathment channels. Lo reporters, but not reporters of liquid-disordered (Ld) membranes, became progressively enriched at axon-associated membrane domains as autotypic junctions matured at ensheathment channels. In the absence of axons, cadherin-enriched lipid microdomains still formed on basal cell membranes, but were not organized into the contiguous domains normally associated with axons. Instead, these isolated domains formed ectopic heterotypic junctions with overlying periderm cells, a distinct epithelial cell type in the epidermis. Thus, axons inhibit the formation of epithelial heterotypic junctions by recruiting cadherin-rich lipid microdomains to form autotypic junctions at ensheathment channels. These findings demonstrate that sensory nerve endings dramatically remodel polarized epithelial components and regulate the adhesive properties of the epidermis.

## Introduction

Epithelia line all of our organs, cavities, and body surfaces, regulating exchange with the outside environment and serving as protective barriers. To carry out these functions, epithelial cells must build polarized membrane domains facing the outside (apical) and inside (basal) environments. Distinct lipids and proteins segregate to apical and basal-lateral membranes, making them biochemically distinct and promoting the formation of polarized structures, including cytoskeletal protrusions and junctions, which join cells into coherent sheets. Junctions also provide instructive polarity cues, enable the generation of specific tissue shapes, impart mechanical strength to tissues, and initiate signaling that responds to the biochemical and mechanical state of the epithelium. Although often overlooked, most epithelia are profusely innervated by sensory or sympathetic neurons. Little work has addressed how neurite branching among epithelial cells affects the polarized organization of epithelial components.

The outer epithelial layer of the skin, the epidermis, serves both as a barrier and sensory organ. Afferent neurites of sensory neurons innervate the epidermis to mediate nociception--the detection of painful mechanical, chemical, or thermal stimuli (Zylka et al., 2005). In mammals, the “free” axonal nerve endings of pain-sensing trigeminal and dorsal root ganglia neurons branch extensively in the epidermis, forming immense receptive fields (Wu et al., 2012).

Although free nerve endings do not contact specialized skin structures (Zylka et al., 2005), recent evidence indicates that these axon endings are functionally coupled (Sondersorg et al., 2014; Baumbauer et al., 2015; Pang et al., 2015; Moehring et al., 2018): Chemo- or optogenetic activation of keratinocytes promotes sensory afferent firing and nocifensive behaviors, while electrically silencing keratinocytes attenuates these behaviors. It has even been suggested that keratinocytes and free nerve endings form synapse-like contacts with one another (Talagas et al., 2020b). Investigating how free nerve endings initiate and maintain contact with keratinocytes is thus essential for understanding the development of the epidermis and its pain-sensing functions.

While much is known about skin-derived signals that attract and pattern peripheral axon arbors (Wang et al., 2013), less attention has been paid to how neurites interact with skin cells once in the epidermis. In both invertebrates and vertebrates, a subset of cutaneous sensory neurons become enveloped by epithelial cell membranes into ensheathment channels, which may be sealed by autotypic epithelial junctions (O’Brien et al., 2012; Tenenbaum et al., 2017; Jiang et al., 2019; Talagas et al., 2020a) or simply engulfed by epidermal membranes(Chalfie and Sulston, 1981). Epidermal ensheathment regulates neurite branching (Tenebaum et al, 2017; Jiang et al, 2019), determines neurite arbor spacing in the skin (Han et al., 2012; Kim et al., 2012; Yang et al., 2019), protects neurites from physical damage (Coakley et al., 2020), and is required for efficient sensory function (Jiang et al., 2019). In flies and zebrafish, the earliest known step in epidermal ensheathment is the accumulation of membranes enriched in a biosensor for the phosphoinositide PI(4,5)P2 (PIP2) at the neurite-skin cell interface (Jiang et al., 2019), indicating that this membrane region either has a distinct lipid composition or greater membrane density. These regions then recruit F-actin in a Rho1-dependent manner (Jiang et al., 2019), presumably to promote curvature of the epidermal membrane to enwrap the neurite. Finally, junction proteins are recruited to seal the neurites within ensheathment channels (O’Brien et al., 2012; Jiang et al., 2019). This sequence of changes in epidermal cell membranes requires the presence of neurites (Jiang et al., 2019), indicating that cues from neurites signal to the skin cell to initiate the ensheathment process. Because most epithelial cells contact many sensory neurites, the formation of autotypic junctions at ensheathment channels has the potential to dramatically redistribute membrane, cytoskeletal components, and junctions in epithelial cells, potentially impacting junction-associated signaling and adhesion between them.

In diverse cellular contexts, membrane domains with distinct lipid compositions function as platforms for signaling and cell-cell interactions. For example, cholesterol and sphingolipids are not distributed uniformly in the plasma membrane, but rather are enriched in domains with lower membrane fluidity, owing to tight packing of the sterol rings on cholesterol with long, saturated acyl chains of sphingolipids (Garcia-Parajo et al., 2014). In simplified model membranes, cholesterol and sphingolipids can undergo large-scale phase separation into micron-sized domains called liquid-ordered (Lo) domains, sometimes equated with lipid “rafts”, which are distinct from so-called liquid-disordered (Ld) regions enriched in unsaturated glycerophospholipids. In native plasma membranes, Lo domains are generally thought to function as nanometer-sized, transient structures that constitute discrete signaling sites in the membrane (Garcia-Parajo et al., 2014). Lo membrane microdomains influence intra- and intercellular signaling by creating platforms for the oligomerization of cell surface receptors and recruitment of downstream effectors (Zuidscherwoude et al., 2014). Lo membranes are enriched at cell-cell junctions (Resnik et al., 2011; Kurrle et al., 2013; Stahley et al., 2014; Lewis et al., 2019), axo-glial junctions (Honke et al., 2002; Schafer et al., 2004), and immune synapses (Dupre et al., 2002; Hiltbold et al., 2003; Nika et al., 2006). Furthermore, the myelin membranes that wrap axons are also highly enriched in cholesterol and sphingolipids, forming phase-separated membrane domains distinct from the rest of the cell membrane (Yurlova et al., 2011). Lo membrane domain enrichment often corresponds to highly curved membranes, including at caveolae (Lamaze et al., 2017), viral budding sites (Sengupta et al., 2019), and filopodia (Scorticati et al., 2011; Honda et al., 2017), where they play instructive roles in membrane remodeling. The observation that axon-associated membranes of epidermal cells are strongly labeled by a PIP2 sensor before neurite ensheathment (Jiang et al., 2019) raises the possibility that membrane microdomains of distinct lipid composition initiate skin cell-neuron interactions. Given the roles of epidermal cells in modulating sensory neuron branching and function, membrane microdomains could be platforms that organize skin cell-neuron interactions during neurite branching in the skin, and in mature skin at sites of functional skin cell-neuron coupling.

To study how sensory axons influence epithelial development, we imaged early stages of epidermal innervation in live zebrafish embryos. During embryonic stages, the peripheral axons of zebrafish somatosensory neurons initially branch between the two epithelial layers of the epidermis: the outer periderm and inner basal layers (O’Brien et al., 2012). Although axons make direct contact with both periderm and basal keratinocytes, as development proceeds they become ensheathed exclusively by the apical membranes of basal cells, and never by periderm cells (O’Brien et al., 2012). Previous work identified the accumulation of a PIP2 biosensor underneath neurites as the earliest known step in neurite ensheathment, promoting the recruitment of downstream ensheathment channel components (Jiang et al., 2019). However, it was unknown whether axon-associated membrane domains (AADs) constitute a unique lipid environment. Here we use quantitative imaging approaches to characterize the maturation of the lipid environment at AADs from the earliest stages of ensheathment to autotypic junction formation. We found that PIP2-containing membrane microdomains are indeed quantitatively enriched in PIP2 and multiple markers for Lo membrane, thus constituting distinct lipid microdomains. PIP2 microdomains are sites of F-actin protrusion formation and preferential cadherin enrichment, well before autotypic junction formation, suggesting that epithelial cells form lipid microdomains as precursors for cell-cell interactions. Consistent with this idea, we found that lipid microdomains mediate keratinocyte-layer-specific interactions with axons.

Moreover, live imaging of the initiation of ensheathment revealed that axons rapidly remodel lipid microdomains through recruitment and de novo formation. In the absence of axons, these cadherin-enriched lipid microdomains formed excessive heterotypic junctions between periderm and basal cells, likely altering the adhesive properties of the epidermis. These findings demonstrate that sensory axons have the potential to dramatically influence basic epithelial cell properties.

## Results

### Axon outgrowth organizes PIP2-containing membrane microdomains on basal keratinocytes

In zebrafish embryos, somatosensory axons initially grow and branch exclusively between the periderm and basal cell layers of the epidermis during the first day of development, and become progressively ensheathed by basal cells over the next several days (O’Brien et al., 2012). To determine how axon outgrowth affects the membranes of periderm and basal epithelial cells, we used confocal live-imaging to follow keratinocyte membrane dynamics from 24-33 hours post-fertilization (hpf) (1 day post-fertilization, dpf, hereafter), a period of active axon branching in the epidermis. To label basal cell membranes, we expressed a UAS-EGFP-PLCγ-PH reporter for the phosphoinositide PIP2 with a *p63*:GAL4VP16 BAC driver line (Rasmussen et al., 2015). Axons were labeled using either a pan-neuronal *neural beta tubulin:DsRed* transgene (Peri and Nüsslein-Volhard, 2008) or a somatosensory-specific *isl1:Gal4;UAS:DsRed* line(Sagasti et al., 2005). As reported previously, EGFP-PLCγ-PH in basal cells initially formed discrete microdomains at the apical surface (Fig 1A, Movie 1). As axons grew over basal cells, EGFP-PLCγ-PH microdomains became more prominent and accumulated underneath axons (Fig 1, A1-A3; Movie 1). These microdomains accumulated immediately upon axon contact, under extending growth cones (Fig 1, A1-3, cyan arrowhead; Movie 1), as well as under axon shafts that had previously grown over basal cells (Fig 1, A1-3, red arrow; Movie 1). These microdomains were highly dynamic and reversibly associated with outgrowing axons (Movie 2). By contrast, axons had dramatically fewer, and qualitatively distinct, effects on periderm membranes labeled with the same reporter. In some cells, periderm membranes were not appreciably affected by axon contact at 1 dpf (Fig 1B, B1-3, cyan arrowheads; Movie 3). In others, outgrowing axons associated with scattered microdomains, but these microdomains did not coalesce into larger axon-associated membrane domains during the imaging interval (Supplemental Fig 1A; Movie 4). To determine the consequence of periderm membrane-axon interactions, periderm cells expressing EGFP-PLCγ-PH and axons were imaged at 4 dpf, a stage when axons are ensheathed by underlying basal cells (O’Brien et al., 2012). We observed two distinct patterns in these periderm cells: either EGFP-PLCγ-PH showed no distinct organization above axons, or it appeared as discrete punctae ordered along axons (Supplemental Fig 1B). Thus, although axons dramatically remodel PIP2-containing microdomains in basal cells, in periderm cells, if such domains form at all, they are discontinuous, indicating that axons preferentially associate with basal cells. Since only basal cells wrap axons into ensheathment channels, early membrane dynamics in basal cells may presage the formation of ensheathment channels (Jiang et al., 2019).

**Figure 1.**
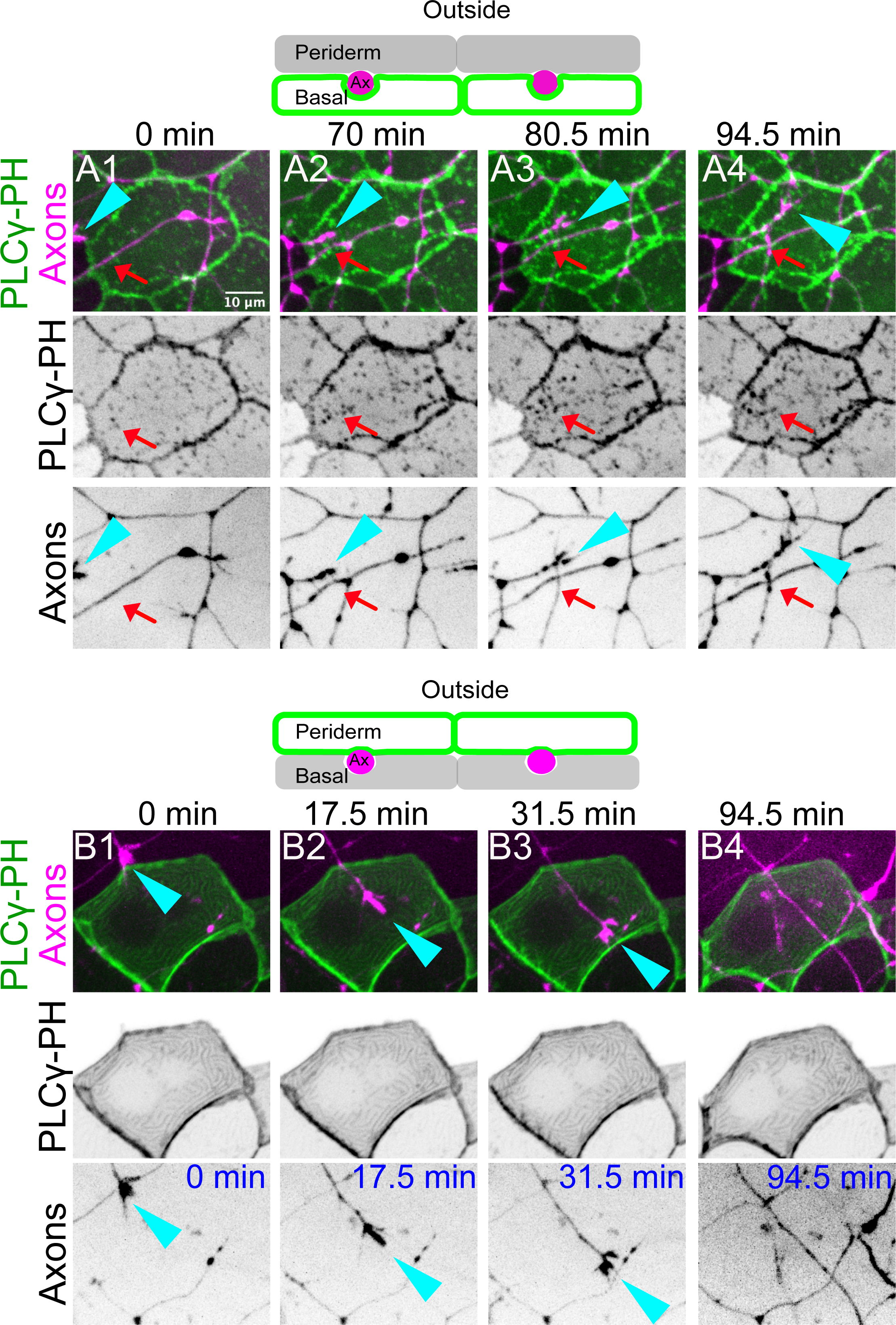
Somatosensory axons induce epidermal-layer specific membrane remodeling. (A) (Top) Schematic of reporter expression in panels A1-A4. The basal cell plasma membrane was labeled with a PIP2 reporter (Tg(*p63:*GAL4VP16; UAS:EGFP-PLCγ-PH, green) and somatosensory neurons (magenta) were labeled with *neural beta tubulin::DsRed* or *isl1:*GAL4VP16; UAS:DsRed transgenes. (A1-A4) Stills from a time-lapse movie. An extending neurite (cyan arrowhead) grew over the apical surface of a basal cell. Shortly after the appearance of the axon, EGFP-PLCγ-PH-labeled membrane microdomains accumulated underneath the axon. Red arrow shows scattered microdomains that accumulated under an axon that had already grown over the basal cell. See Movie 1. (B) (Top) Schematic of reporter expression in panels B1-B4. Periderm cell plasma membranes (green) were mosaically labeled with a UAS:EGFP-PLCγ-PH transgene injected into Tg(*krt5:*GAL4FF); Tg(*neural beta-tubulin:*DsRed) embryos. Axons are labeled in magenta. (B1-B4) Stills from a time-lapse movie. Axons had no apparent effect on the plasma membrane of overlying periderm cells. Cyan arrowhead marks a growth cone crawling underneath the labeled periderm cell. See Movie 3.

### Basal cell microdomains have a distinct lipid composition

Axon-associated microdomains in neurite-ensheathing epithelial cells in both flies and fish are enriched in EGFP-PLCγ-PH (Jiang et al., 2019). These regions of increased fluorescence could be quantitatively enriched in the PIP2 lipid, or simply contain higher membrane densities. To distinguish between these possibilities, we generated a dual reporter that expressed the EGFP-PLCγ-PH sensor bicistronically with mRuby-CAAX, a potentially “generic” prenylated membrane reporter, using a self-cleaving T2A peptide. This dual membrane reporter was placed under the control of the multimerized Upstream Activating Sequence (UAS:EGFP-PLCγ-PH-T2A-mRuby-CAAX) and injected into *p63:GAL4VP16* transgenic animals to mosaically label basal cells. Embryos were imaged by confocal microscopy at 1 dpf, when microdomains begin accumulating underneath axons. To estimate the levels of each reporter recruited into microdomains, we measured the ratio of each reporter inside versus outside microdomains (see Materials and Methods), providing a measure of relative reporter enrichment. EGFP-PLCγ-PH and mRuby-CAAX both labeled membrane microdomains in basal keratinocytes (Fig 2B, cyan arrowheads) and had relative enrichment values >1 (Fig 2B), indicating that membrane microdomains are most likely regions of increased membrane surface area. However, EGFP-PLCγ-PH was consistently more enriched in these domains than mRuby-CAAX (Fig 2C). Thus, membrane microdomains strongly labeled by EGFP-PLCγ-PH are indeed enriched in PIP2, defining areas of distinct lipid composition in the apical membrane of basal keratinocytes.

**Figure 2.**
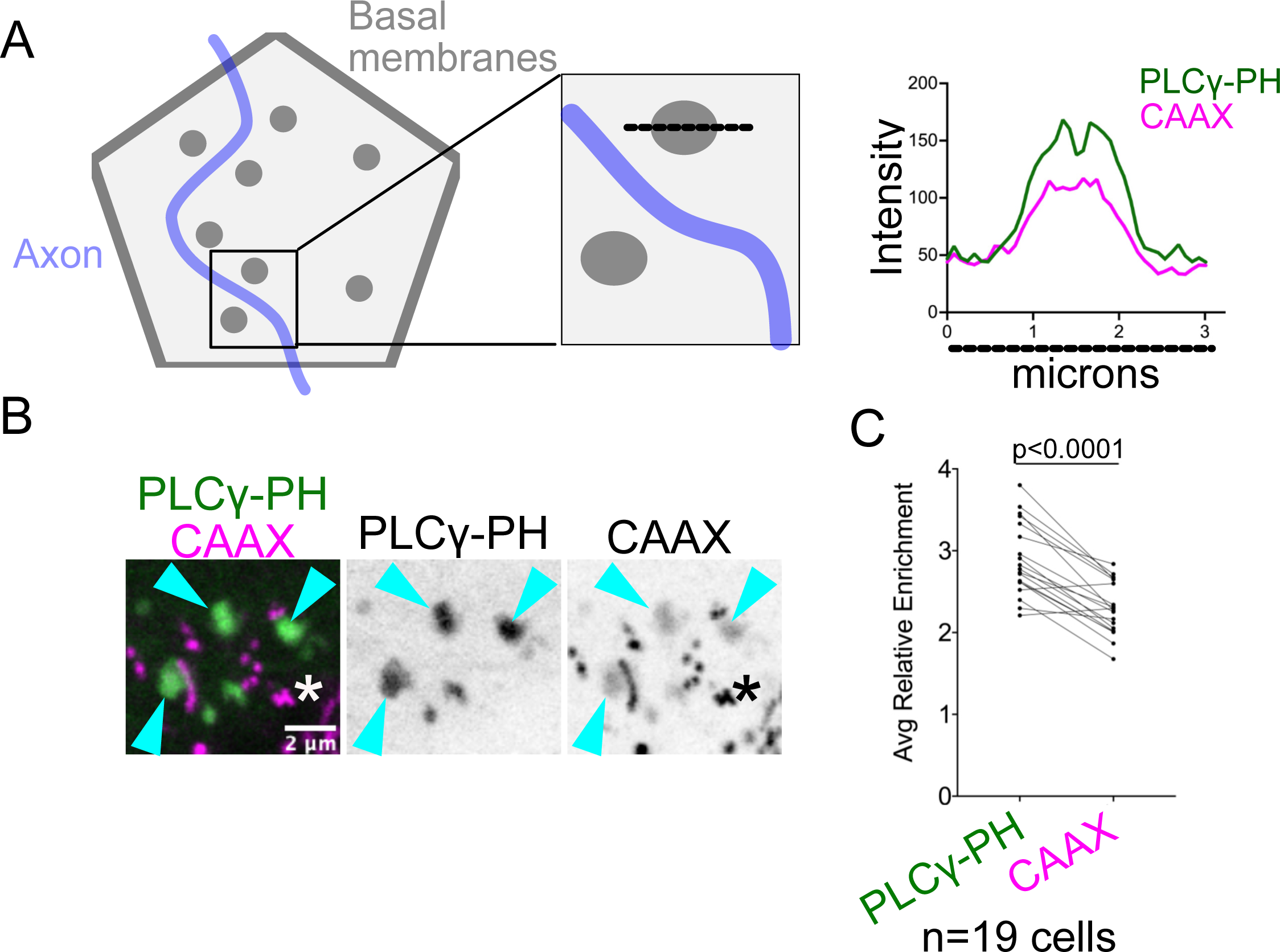
Basal cell membrane microdomains are enriched in a PIP2 reporter. (A) (Left) Schematic showing how relative reporter enrichment was quantified. Lipid microdomains are represented in gray; Axon (not imaged) in blue. Intensity plots for EGFP-PLCγ-PH and mRubyCAAX were generated along a line drawn across microdomains (dotted black line). (Right) Intensity plot from a representative microdomain. (B1-3) Image of basal cell membrane co-expressing EGFP-PLCγ-PH and mRubyCAAX at 1 dpf. Microdomains were labeled by both reporters, but GFP was more enriched. mRubyCAAX also accumulated in numerous aggregates and/or endosomes in basal cells (asterisk). (C) Quantification of relative enrichment of the two membrane reporters in isolated microdomains (204 microdomains measured from 19 cells from 7 embryos). Enrichment values from the same basal cell clones were averaged to obtain the average relative enrichment for each cell. EGFP-PLCγ-PH was significantly more enriched in microdomains than mRubyCAAX (p<0.0001, paired t-test).

### Axon-associated PIP2 microdomains form both by reorganizing existing domains and forming them *de novo*

PIP2 lipid microdomains could accumulate underneath axons either through the lateral coalescence of pre-formed microdomains or *de novo* formation underneath axons. To determine how axons accumulate lipid microdomains in the underlying basal cell membrane, we made time-lapse confocal movies of these microdomains with higher magnification, and found evidence for both mechanisms.

During lateral coalescence, microdomains became ordered along filopodial tips (Fig 3A2; Movie 5) before accumulating under the axon shaft proximal to the growth cone (Fig 3A3-5; Movie 5). If axons promote the lateral coalescence of lipid microdomains on the basal cell surface, we predicted that microdomains would remain scattered in the absence of cutaneous axons. To test this prediction, we made time-lapse movies of basal cells in *neurogenin-1* (*ngn1)* morphants that entirely lack somatosensory skin innervation (Andermann et al., 2002; O’Brien et al., 2012). In *ngn1* morphants, basal cells formed PIP2 lipid microdomains, but these domains did not coalesce into extended structures over the course of 3 hours at 1 dpf (Supplemental Fig 2A; Movie 6). Thus, axons are not required for microdomain formation, but rather to direct their reorganization.

**Figure 3.**
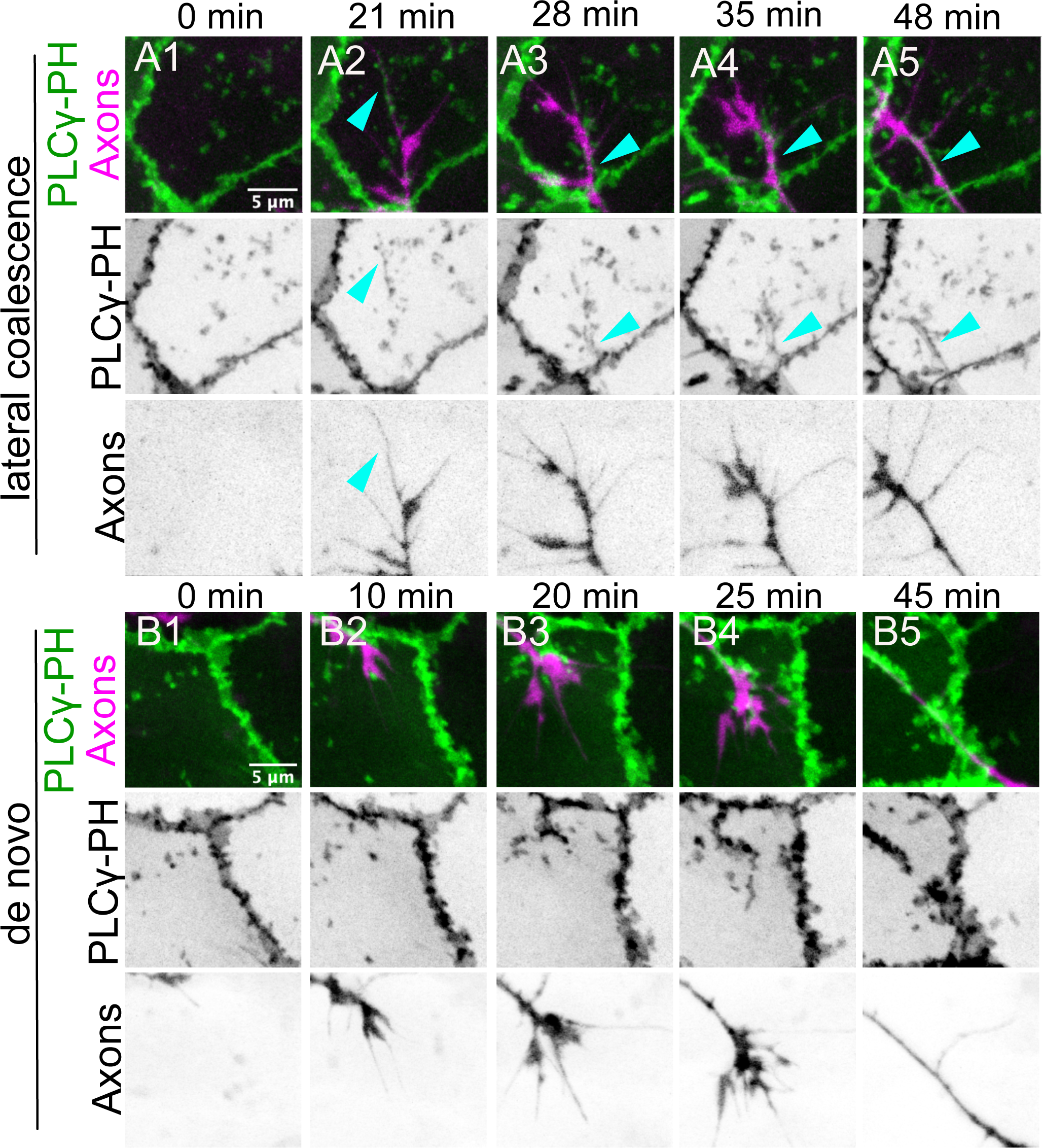
Axons reorganize epithelial PIP2 microdomains by lateral coalescence or de novo formation. (A1-A5) Stills extracted from a timelapse movie of a Tg(*p63-*GAL4-VP16:UAS:EGFP-PLCγ-PH); Tg(*isl1:LexA:LexAop:TdTomato*) 1 dpf embryo (3 min per frame). (A1) Prior to the appearance of a neurite, the basal cell surface contained isolated PIP2-rich microdomains. (A2) Below an advancing growth cone, microdomains ordered along filopodial tips (cyan arrow). (A3) Lipid microdomains coalesced underneath an axon shaft proximal to the growth cone (cyan arrow), forming an axon-associated domain (AAD). (A4-5) An AAD elongated along an extending axon. See Movie 5. (B1-5) Timelapse of axons in a Tg(*p63-*GAL4-VP16: UAS:EGFP-PLCγ-PH); Tg(*neural beta tubulin::DsRed*) 1 dpf embryo (5 min per frame). Shortly after a growth cone passed over the pictured basal cell, lipid microdomains formed directly below the growth cone and associated with the elongating axon shaft. See Movie 7.

We also found evidence for *de novo* formation of axon-associated lipid microdomains. These domains formed rapidly after an axon passed over the apical surface of a basal cell otherwise sparsely containing lipid microdomains (Fig 3B1-5; Movie 7). A potential mechanism of *de novo* microdomain formation was suggested while capturing basal cell division events during axon outgrowth (Movie 8). Rounding mitotic basal cells pulled on nearby non-dividing basal cells; PIP2 microdomains rapidly disassembled as the non-dividing cell was pulled toward the dividing cell. As the non-dividing cell relaxed, microdomains formed rapidly and specifically around axon shafts (Movie 8). This observation suggests that microdomain formation could involve a local reduction in membrane tension as an axon presses against the basal cell membrane. *De novo* formation could, therefore, reflect a passive remodeling of the basal cell surface as axons crawl over them (e.g. depression). However, lipid microdomain accumulation is restricted to some axonal segments, while others induce no remodeling of the basal cell surface (Supplemental Figure 2B, Movie 9), suggesting that microdomain accumulation is a consequence of specific recognition of axons, perhaps those destined to be ensheathed by basal cells.

### Lipid microdomains are sites of F-actin-based protrusion formation

PIP2 can regulate the cortical actin cytoskeleton by recruiting several actin-binding proteins (Janmey et al., 2018). To determine if F-actin associates with basal cell PIP2 microdomains during axon outgrowth, we imaged the actin reporter LifeAct-mRuby. To co-visualize lipid microdomains and the actin cytoskeleton, the UAS:LifeAct-mRuby plasmid was injected into *p63*:GAL4VP16;UAS:EGFP-PLCγ-PH eggs, and embryos were imaged at 1 dpf by time-lapse confocal microscopy. Isolated PIP2 lipid microdomains were enriched in LifeAct signal (Fig 4A and Fig 4B, Movie 10), demonstrating that lipid microdomains define sites of preferential F-actin polymerization and/or stabilization. However, F-actin was not recruited uniformly at lipid microdomains, but rather was concentrated within subregions (Fig 4B1 and Fig 4B2), suggesting that lipid microdomains are compartmentalized. LifeAct-mRuby fluorescence at lipid microdomains fluctuated over time as individual lipid microdomains remodeled (compare Fig 4B1 and 4B2) and sometimes receded while the PIP2 microdomain remained stable (Fig 4B4). Thus, PIP2 lipid microdomains are sites of dynamic F-actin remodeling, but PIP2 enrichment alone is not sufficient for F-actin stabilization. We propose that PIP2 lipid microdomains may reversibly recruit factors that promote the stability of F-actin at microdomains.

**Figure 4.**
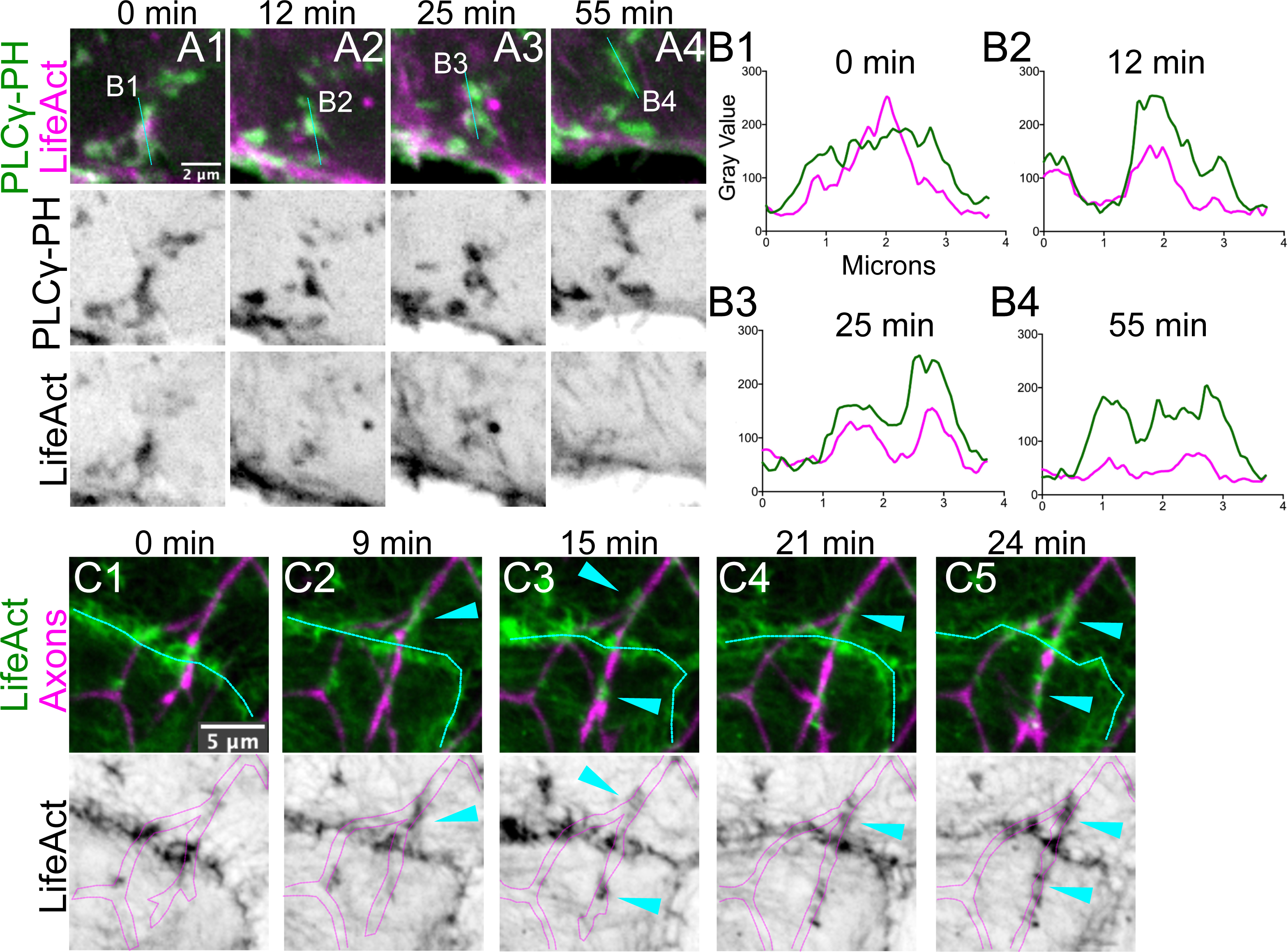
Basal cells form F-actin protrusions around axons. (A1-A4) Stills from movies of 1 dpf Tg(*p63:GAL4VP16;*UAS:EGFP-PLCγ-PH) injected with a 10xUAS:mRuby-LifeAct plasmid. The F-actin reporter signal reorganized concomitantly with microdomain remodeling. See Movie 10. (B1-B4) Plots showing fluorescence intensity profiles along the dotted cyan lines in A1-A4, showing colocalization of increased LifeAct signal with PIP2 microdomains. F-actin polymerized/stabilized transiently at PIP2 microdomains (compare panels B1-3 with B4). (C1-4) Stills from movie of 1 dpf Tg(*p63:GAL4VP16;UAS:LifeAct-GFP*); Tg(*neural beta tubulin::DsRed*) fish (3 min intervals). Filamentous LifeAct-GFP signal dynamically wrapped axonal segments. Protrusions with LifeAct-GFP signal completely surrounded the axon. Dotted line marks boundary between basal cells. See Movie 12.

The accumulation of F-actin at microdomains suggests that these domains could form actin protrusions that associate with axons. To test this hypothesis, we made time-lapse movies of cutaneous axons growing over basal cells expressing LifeAct-GFP (*p63*:GAL4VP16;UAS:LifeAct-GFP;(Rasmussen et al., 2015). While extending growth cones induced no observable effect on basal cell F-actin organization (Movie 11), protrusive actin structures dynamically wrapped around axon shafts (Fig 4C cyan arrowheads; Movie 12). In some cases, basal cells formed protrusions parallel to the apical surface, allowing their structure to be unambiguously discerned (Movie 13): in those instances, EGFP-PLCγ-PH fluorescence appeared concomitantly with F-actin appearance, suggesting an association between PIP2 production and F-actin polymerization at the apical surface of basal cells. F-actin associated with axon shafts underwent continuous remodeling (Fig 4C2-5, cyan arrowheads). As with EGFP-PLCγ-PH, F-actin association with axons was selective, with some axon branches eliciting numerous F-actin protrusions (Movie 14, cyan arrowhead) whereas others were ignored by basal cells (Movie 14, red arrowhead). We conclude that PIP2 lipid microdomains are sites of dynamic F-actin polymerization/stabilization, resulting in the formation of membrane protrusions that reversibly wrap axons. F-actin production may allow protrusions to extend and retract dynamically during this process to sample the axonal surface and distinguish axons that will be ensheathed from those that will not.

### Axon-associated microdomains become progressively enriched in liquid-ordered reporters

Since axon-associated microdomains have a distinct lipid composition (Fig 2), we asked whether axon-associated membranes had other distinct lipid properties. Lipid rafts are membrane “nanodomains” enriched in cholesterol and sphingolipids; the bulky hydrophobic moieties in these lipids promote the formation of nanometer-sized liquid-ordered (Lo) membrane phases (Fig 5A) that can coalesce into microdomains that serve as platforms for cell-cell interactions (Zuidscherwoude et al., 2014). To determine if axon-associated membrane domains are enriched in Lo membranes, we generated dual membrane reporters based on previously validated reporters that quantitatively partition into Lo and Ld (raft vs non-raft) membranes (Fig 5A) (Pyenta et al., 2001; Zacharias et al., 2002; Sengupta et al., 2019). We used the double myristoylated and palmitoylated lipid modifications (mp-mEGFP and mp-mApple) and a GPI-anchored SuperFolder GFP (sfGFP-GPI) as Lo reporters, and adapted geranylgeranylated fluorescent reporters (mEGFP-gg and mApple-gg) as Ld reporters. Because intrinsic fluorescent protein properties can promote Lo-membrane-independent clustering (Zacharias et al., 2002), we used a variant of EGFP mutated to prevent oligomerization (mp-EGFP A206K, L221K, F223K, hereafter called mp-mEGFP; see Materials and Methods). We co-expressed Lo and Ld reporters in the same basal cells using a T2A peptide to measure the relative enrichment of each reporter within (“In”) and outside (“Out”) axon-associated membranes (Fig 5B).

**Figure 5.**
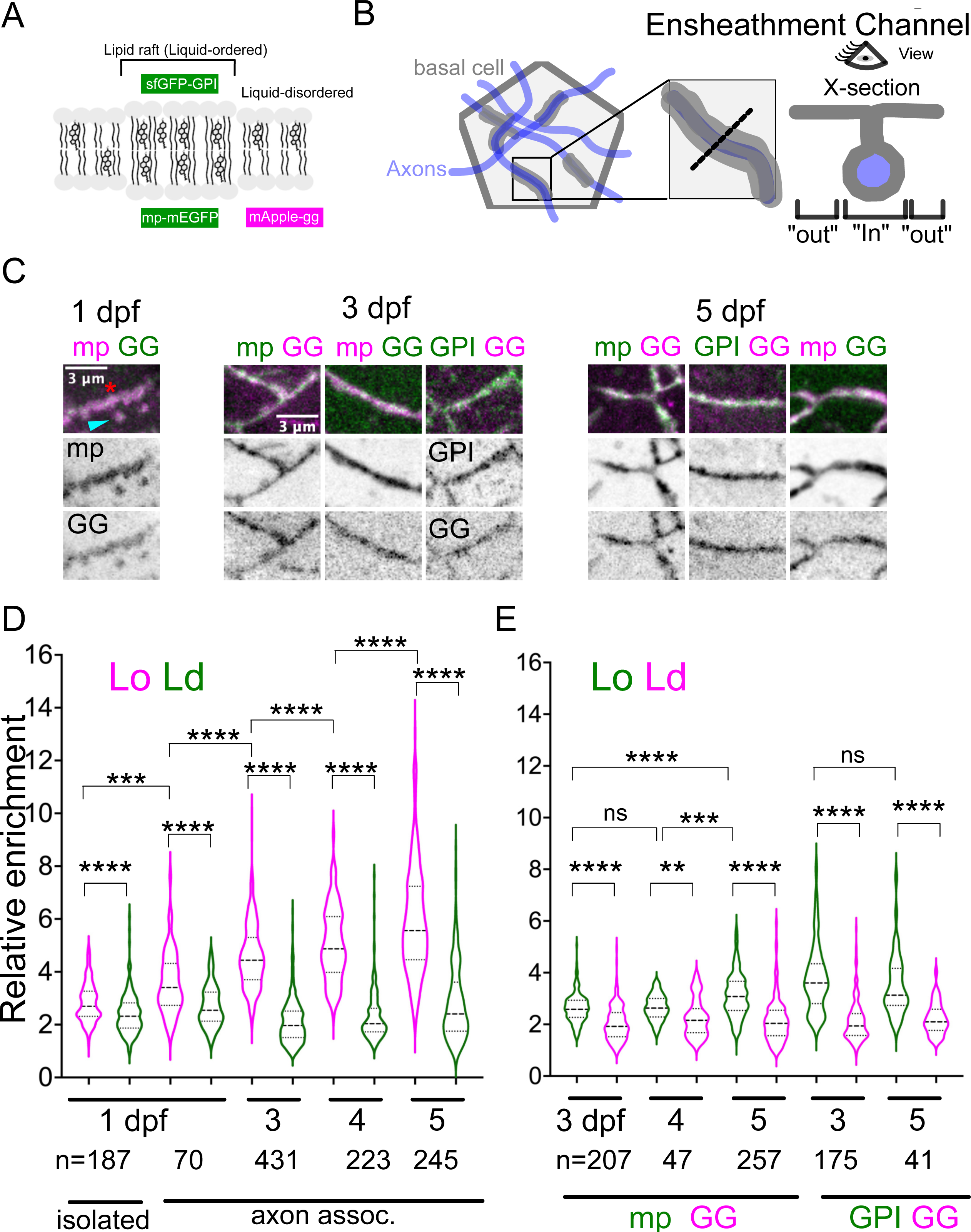
Axon-associated domains progressively enrich reporters for Lo, but not Ld, membrane. (A) Double myristoylation and palmitoylation, or modification by glycosylphosphatidylinositiol anchors, localizes fluorescent protein reporters preferentially in Lo membrane. Gernanylgeranylated fluorescent reporters were used to label Ld membrane. (B) Schematic of membrane reporter localization (gray) with respect to axons (blue). Inset shows detail of axon-associated membrane domain (AAD). Relative enrichment of each membrane reporter was measured using the ratio of each reporter inside (“In”) to outside (“Out”) AADs. (C) Details of AADs in basal cells expressing Lo and Ld reporters bicistronically. Lo and Ld reporters were imaged between 1-5 dpf. At 1 dpf, relative enrichment was measured in both isolated microdomains (cyan arrowhead in C; “isolated” in D) and larger axon-associated domains (red asterisk in C) (D) Relative enrichment of AADs from basal cells expressing mp-mApple-T2A-mEGFP-gg. mp-mApple (Lo reporter) exhibited significantly greater relative enrichment than mEGFPgg (Ld reporter) at 1, 3, 4, and 5 dpf (**** p<0.0001, Wilcoxon signed rank tests). Over time, the relative enrichment of mp-mApple increased significantly (*** p<0.001, ANOVA with paired t-test; **** p<0.0001, ANOVA with Mann-Whitney U test). (E) Relative enrichment of AADs from basal cells expressing mp-mEGFP-T2A-mApple-gg or sfGFP-GPI-T2A-mApple-gg at 3, 4, and 5 dpf. Greater relative Lo enrichment was independent of FP combination, and can be recapitulated using an independent Lo reporter (sfGFP-GPI) (** p<0.01, **** p<0.0001, Wilcoxon signed rank tests). mp-mEGFP shows significantly greater enrichment at 5 dpf versus 4 dpf (*** p<0.001, ANOVA with Mann-Whitney U test) and versus 3 dpf (**** p<0.0001, ANOVA with Mann Whitney U test).

Isolated microdomains at 1 dpf were more enriched in the Lo reporter mp-mApple than the Ld reporter mEGFP-gg (Fig 5C, D; p=7.21e -11, Wilcoxon signed rank test). Increased relative enrichment of the Lo reporter persisted following the formation of larger axon-associated microdomains and mature ensheathment channels, from 1-5 dpf (Fig 5D,1 dpf :p=8.31e-7; 3-5 dpf: p<2.2e-16; Wilcoxon signed rank tests; Supplemental Fig 3A). To rule out fluorescent protein-specific effects, we swapped the fluorophores between the dual Lo/Ld reporters (mp-EGFP-T2A-mApple-gg). As with mp-mApple, mp-mEGFP was more enriched than the Ld reporter between 3-5 dpf (Fig 5E, 3 dpf: p<2.2e-16; 4 dpf: p=0.002; 5 dpf: p<2.2e-16; Wilcoxon signed rank tests; Supplemental Fig 3B). To rule out the possibility that the putative Lo enrichment was an artifact of the myristoylation and palmitoylation modifications, we used the sfGFP-GPI reporter for Lo membranes. Similar to mp-mApple and mp-mEGFP, sfGFP-GPI was significantly more enriched at axon-associated domains than mApple-gg (Fig 5E, 3 dpf: p < 2.2e-16; 5 dpf: p=6.23e-11; Wilcoxon signed rank tests; Supplemental Fig 3C). Lower relative enrichment of Ld reporters (mAppleGG and mEGFPGG) was at least partly due to increased cytosolic signal compared to the Lo reporters, which showed more stable membrane recruitment, confirming that the different lipid modifications used in these reporters are distinctly regulated. We also found that EGFP-PLCγ-PH exhibited greater relative enrichment at axon-associated membranes compared to mApple-gg (Supplemental Figure 3D,E), suggesting that PIP2 may also be enriched within the Lo membranes around axons. Collectively, these results indicate that distinct lipid modified proteins show specific preferences for axon-associated membranes, raising the possibility that lipid-raft-enriched membranes accumulate preferentially around axons.

Myristoylated and palmitoylated Lo reporters were increasingly enriched at axon-associated membrane domains from 1-5 dpf, whereas relative enrichment of the Ld reporter remained relatively constant (Fig 5D and E; see Supplemental Table 1 for statistics). This observation suggested that axon-associated membranes mature over time by changing their lipid composition. The GPI-anchored Lo reporter showed the greatest relative enrichment regardless of stage, indicating that sfGFP-GPI recruitment reaches its maximum earlier than the myrpalm Lo reporter. Collectively, these results suggest that lipid microdomain association with axons promotes the accumulation of Lo membranes at sites of future ensheathment channels, and could facilitate ensheathment channel formation by clustering proteins and/or promoting membrane curvature (Sengupta et al., 2019).

### Lipid microdomains accumulate junction proteins that contribute to ensheathment channel formation

The final step of sensory neurite ensheathment is the recruitment of junctions to epithelial membranes surrounding neurites (Jiang et al., 2019). Since junction-forming proteins can partition preferentially into Lo membranes (Lewis et al, 2019), we hypothesized that lipid microdomains in basal cells enrich cadherins that will later form autotypic junctions at ensheathment channels. To test this idea, we expressed fluorescently tagged adherens junction and desmosomal cadherins, which are both found at ensheathment channels (Jiang et al., 2019). To visualize E-cadherin, we used a *cdh1-TdTomato* gene trap line reporting endogenous E-cadherin (E-cad) localization (Cronan et al., 2018). To visualize Desmocollin 2-like (Dsc2l) we used a bacterial artificial chromosome (BAC) reporter with GFP-tagged Dsc2l. Isolated PIP2 lipid microdomains in basal cells were slightly enriched in E-cad (Fig 6A, B; left C) and strongly enriched in Dsc2l (Fig 6A, B; left D) at 1 dpf, well before ensheathment channels form. Cadherin localization to microdomains was heterogeneous, often appearing to cluster within subregions of PIP2 microdomains (Fig 6B), suggesting that their enrichment to microdomains was not simply a consequence of increased membrane in those areas. Interestingly, Dsc2l showed stronger relative enrichment than E-cadherin at microdomains (compare Fig 6C and 6D), suggesting that desmosomal and adherens junction cadherins may be recruited or stabilized at lipid microdomains by distinct mechanisms. Thus, microdomain accumulation at axon contact sites may promote the recruitment of junction proteins needed for subsequent ensheathment.

**Figure 6.**
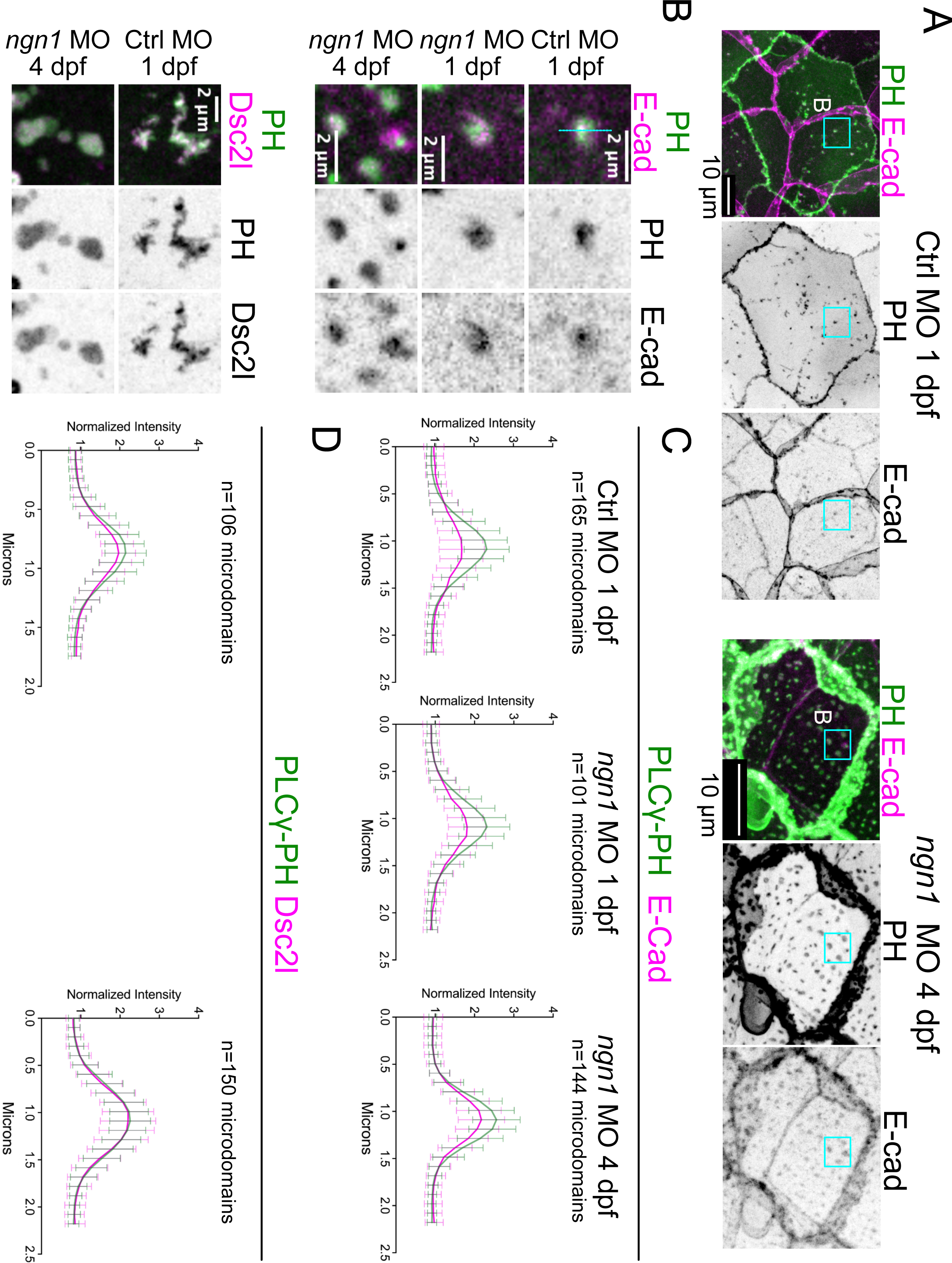
Lipid microdomains accumulate Cadherins independently of axons. (A) To measure E-Cadherin recruitment to lipid microdomain prior to axon association, Tg(*p63:GAL4VP16*; UAS:EGFP-PLCγ-PH); *cdh1-*TdTomato embryos were injected with a control morpholino (Ctrl MO) and imaged at 1 dpf. To measure E-Cadherin recruitment in the prolonged absence of axons, Tg(*p63:GAL4VP16*; UAS:EGFP-PLCγ-PH); *cdh1-*TdTomato embryos were injected with a *neurogenin-1* morpholino (*ngn1* MO) and imaged at 4 dpf. Shown are representative images from the E-Cadherin experiments. (B) Representative images of E-cadherin (Top) and Dsc2l (bottom) recruitment to lipid microdomains prior to axon association in wild-type control embryos (Ctrl MO 1 dpf), *ngn1* MO embryos (*ngn1* MO 1 dpf), and in the prolonged absence of cutaneous axons (*ngn1* MO 4 dpf). Prior to association with axons (Ctrl MO 1 dpf), PIP2 lipid microdomains showed a slight enrichment of E-Cadherin and a stronger enrichment in Dsc2l. E-Cadherin recruitment to microdomains was axon-independent (*ngn1* MO 1 dpf). In the prolonged absence of cutaneous axons (*ngn1* MO 4 dpf), E-cadherin recruitment increased and Dsc2l enrichment was similar to that of EGFP-PLCγ-PH.

To test if microdomain lipid composition promotes the recruitment of junction proteins, we treated fish acutely with methyl-beta-cyclodextrin (MBCD), which disrupts Lo membranes by extracting cholesterol (Zidovetzki and Levitan, 2007). This treatment modestly reduced E-cadherin enrichment at ensheathment channels (Supplemental Figure 3F). While the effects of MBCD are not restricted to Lo membrane domains (Zidovetzki and Levitan, 2007), this result suggests that Lo membranes may be required for the maintenance of junction proteins at ensheathment channels.

### Lipid microdomains enrich E-cadherin independently of axons

Basal cell cadherins may be recruited to ensheathment channels by axons directly, or by PIP2- and Lo-enriched microdomains that coalesce into axon-associated domains. To distinguish between these possibilities, we asked if cutaneous axons are required for cadherin recruitment to microdomains, and if PIP2 lipid microdomains in basal cells mature independently of axons. To test the requirement for axons in cadherin recruitment to lipid microdomains, we compared endogenous E-cad recruitment in microdomains in 1 dpf control morphants, which had not yet coalesced into axon-associated domains, to its recruitment in microdomains in *ngn1* morphants. E-cad was enriched to a similar extent in wild-type control and *ngn1* micromains at 1 dpf (Fig 6C, left and middle). To determine whether lipid microdomains continue to mature in the prolonged absence of axons, we measured E-cad enrichment in *ngn1* morphants at 4 dpf, when axons are normally sealed by autotypic junctions into ensheathment channels. Compared to “early” microdomains in 1 dpf control morphants, “late” microdomains in *ngn1* morphants were more enriched in E-cad (Fig 6B,C). Early and late microdomains exhibited equivalent recruitment of Dsc2l (Fig 6B, D). Thus, axons are not required for cadherin recruitment to microdomains, indicating that lipid microdomains mature independently of ensheathment channel formation.

### Axon-mediated recruitment of Cadherin-enriched microdomains into ensheathment channels inhibits the formation of basal cell-periderm heterotypic junctions

What happens to lipid microdomains in the absence of axons? The observation that they continue to accumulate cadherins suggested that the absence of axons might cause increased heterotypic junctions to form between periderm and basal cells. This model predicts that, in the absence of axons: 1) The number of apical microdomains in basal cells should increase; 2) these domains should stabilize over time; 3) similar stable domains in periderm membranes should associate with apical microdomains in basal cells; and 4) that increased basal-periderm junctions should be detectable ultrastructurally.

To test the first prediction, we compared the density of isolated PIP2+Ecad+ microdomains in control and *ngn1* morphants at 3 dpf (Fig 7A-C, cyan arrowheads), a stage when axons are sealed by autotypic contacts into ensheathment channels in wildtype animals (red arrows in Fig 7A) (O’Brien et al., 2012). In the control epidermis, PIP2+Ecad+ microdomains were observed primarily at axon ensheathment channels; isolated PIP2+Ecad+ microdomains formed in regions devoid of ensheathment channels (Fig 7A, cyan arrows). Consistent with this prediction, in the absence of axons, the number of isolated PIP2+Ecad+ microdomains on basal cell membranes dramatically increased (Fig 7B,C).

**Figure 7.**
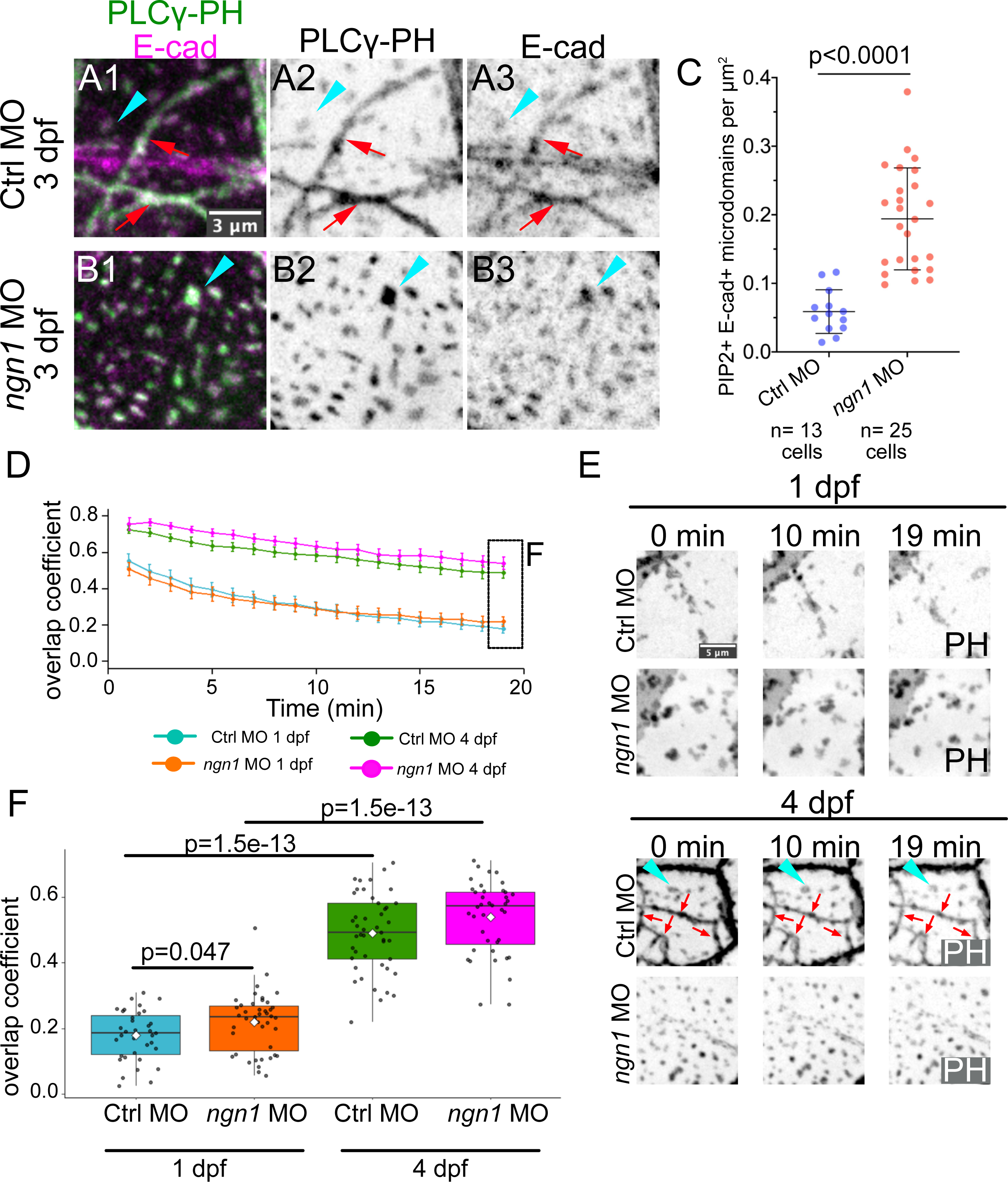
In the absence of axons, microdomains mature into stable periderm-basal contacts. (A) Tg(*p63:*GAL4VP16;UAS:EGFP-PLCγ-PH); *cdh1-*TdTomato embryos injected with control morpholino (Ctrl MO) or *ngn1* MO and imaged at 3 dpf. In wild-type controls (A1-3), E-cadherin in the basal layer localized to ensheathment channels (red arrows) and isolated microdomains (cyan arrowhead). In the absence of axons (B1-3, C), basal cells formed an increased number of E-cadherin-containing isolated microdomains (p value computed using Welch’s t-test, 3 embryos per condition). (D-F) 1-min interval time-lapse movies of basal cell lipid microdomains were made in Tg(p*63:*GAL4VP16*:UAS:*EGFP-PLCγ-PH) embryos injected with control MO or *ngn1* MO. See Movie 15. In (D), the overlap coefficient of each frame to the first frame of the movie is plotted over time to measure the dynamics of lipid microdomains unassociated with axons in 4 conditions. (E) Stills from representative movies are shown. Cyan arrowhead shows an example of a microdomain unassociated with ensheathment channels (red arrows). At 1 dpf, isolated lipid microdomains were highly dynamic structures, while at 4 dpf they were static structures. (F) Comparison of end-point overlap coefficients for individual basal cells for each condition (p values computed using Mann-Whitney U tests. Ctrl MO 1 dpf, n=35 basal cells from 5 embryos; *ngn1* MO 1 dpf, n=43 cells from 4 embryos; control MO 4 dpf, n=45 cells from 4 embryos; *ngn1* MO 4 dpf, n=40 cells from 4 embryos).

To test the second prediction--that microdomains should stabilize over time--we analyzed microdomain dynamics at the apical membrane of basal cells in the presence and absence of axons in Tg(*p63*:GAL4VP16;UAS:EGFP-PLCγ-PH) embryos (Fig 7D-F). To quantify microdomain dynamics, we made 1 min interval movies of EGFP-PLCγ-PH-labeled lipid microdomains and measured the overlap coefficient of each frame to the first frame. At 1 dpf, microdomains in *ngn1* morphants showed comparable dynamics to isolated microdomains in control morphants at the same stage (Fig 7D, cyan and orange lines; Movie 15). However, in 4 dpf *ngn1* morphants, membrane microdomains became markedly static structures, showing significantly higher overlap coefficients than microdomains from 1 dpf *ngn1* morphants (Fig 7D, F; Movie 15). The dynamics of these ectopic lipid microdomains in 4 dpf *ngn1* morphants closely matched those of relatively sparse microdomains not associated with ensheathment channels in wild-type controls at the same stage (Fig 7D, F; Movie 15), indicating that the formation of stable microdomains is not an artifact of the complete loss of axons in the skin. We conclude that, over time, lipid microdomains not associated with axons reduce their lateral motion and become static membrane domains.

Though lipid microdomains mature independently of axons, axons modestly promoted microdomain dynamics in 1 dpf embryos (Fig 7F). While imaging lipid microdomains in 1 dpf WT embryos, we observed instances of transient lipid microdomain ordering directly underneath Cadherin-rich periderm cell membranes--a phenomenon particularly prominent in the absence of axons (Supplemental Fig 4A,B). When microdomains were compared within single basal cells from 1 dpf *ngn1* morphants, ordered microdomains had reduced dynamics compared to scattered ones (Supplemental Fig 4C, D), suggesting that microdomains can be immobilized through homophilic interactions at Cadherin-rich membranes in the absence of axons. Thus, axons promote the remodeling of basal cell surfaces by preventing ectopic association of lipid microdomains with Cadherin-containing periderm membranes.

To test the third prediction--that lipid microdomains in basal cells align with lipid microdomains in periderm cells--we simultaneously labeled Lo membrane in periderm cells using a periderm-specific Tg(*krt5:*mp-mCherry) reporter and PIP2 in basal cells using Tg(*p63:*GAL4VP16; UAS:EGFP-PLCγ-PH). In 3 dpf larvae injected with *ngn1* MO, ∼50-100% of all stable microdomains on the apical surface of basal cells associated with microdomains enriched in mp-mCherry on the basal surface of periderm cells (n= 56 cell pairs from 4 embryos) (Fig 8A,B).

**Figure 8.**
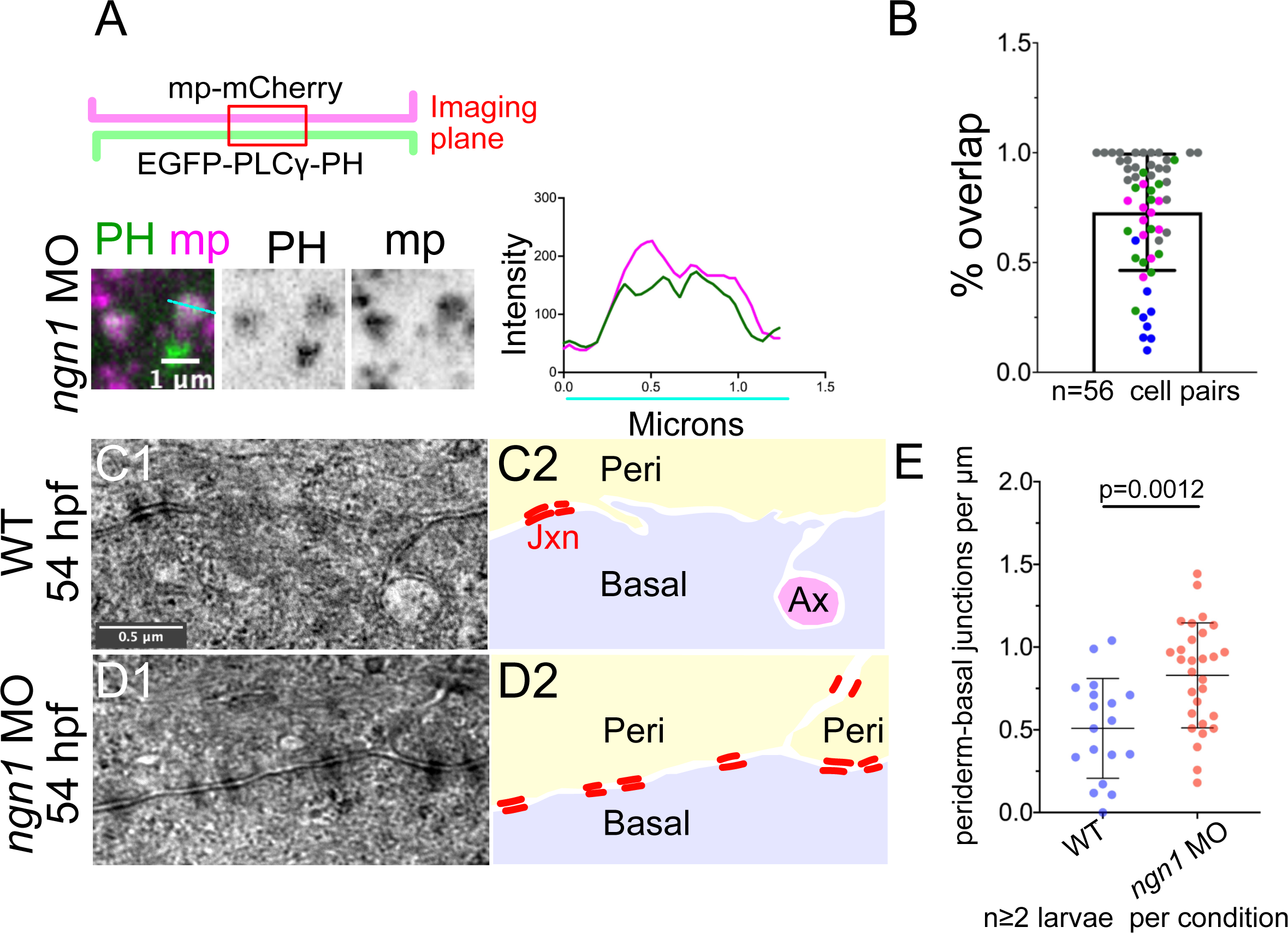
Axons inhibit the formation of periderm-basal cell junctions. (A) (Top) Diagram of membrane labeling in experiments shown below. (Left) Periderm membrane (magenta) was specifically labeled using the krt5 promoter (Tg(*krt5:*mp-mCherry)), and basal membrane was specifically labeled using the *p63:*GAL4VP16 driver (Tg(*p63:*GAL4VP16; UAS:EGFP-PLCγ-PH)) in larvae injected with *ngn1* MO to block sensory neuron development. Larvae were imaged at the interface between periderm (magenta) and basal (green) membranes. Intensity plot shows gray values for green and red channels along the indicated line. (B) Quantification of the proportion of basal cell microdomains that overlapped with periderm microdomains. Each dot (n=56) represents an independent basal-periderm cell pair. Colors indicate data from four different larvae; An average of 24 microdomains were scored per basal cell. (C,D) Representative TEM images of periderm-basal interfaces in 54 hpf wild-type (WT, C1) or *ngn1* morphants (*ngn1* MO, D1). Junctions were identified as electron-dense plaques at the membrane (red bars in schematics on right). In WT epidermis, periderm-basal cell junctions were observed alongside wrapped axons (Ax in schematic). In the absence of axons, the density of ultrastructurally-defined periderm-basal junctions increased (** p<0.0012, unpaired t-test). Each data point represents a unique region of the epidermis from at least 2 larvae per condition.

To directly quantify periderm-basal cell junctions, we analyzed previously generated transmission electron microscopy data sets of WT control and *ngn1* morphants (O’Brien et al., 2012). Periderm-basal cell boundaries were followed in TEM sections in 54 hpf larvae. Junctions were scored as discrete electron-dense plaques on opposing periderm and basal membranes, with intervening electron-density in the intermembrane space. We found an increased density of ultrastructurally defined junctions at periderm-basal boundaries in the absence of cutaneous axons versus WT controls (Fig 8C-E). These data indicate that cutaneous axons remodel epithelial contacts during skin innervation, sequestering junction proteins at ensheathment channels that would otherwise contribute to inter-keratinocyte adhesion. We propose that basal keratinocytes recruit and sequester junction components into lipid microdomains that function as versatile adhesion platforms to maintain the proper balance between axon ensheathment and epithelial adhesion.

## Discussion

Using live imaging of the larval zebrafish epidermis, we have found that cutaneous free axon endings remodel lipids, actin, and junctions in basal keratinocytes. Most studies of axon outgrowth have focused on how growth cones crawl over ECM substrates using integrin-based adhesions, but axons can also crawl over the membranes of other cells. How nascent neurites interact with cellular substrates is poorly understood. Our live imaging experiments demonstrate that epithelial cell membranes are not passive substrates that growth cones crawl over, but rather respond dynamically to axon contacts. We found that axons capture and create membrane microdomains of distinct lipid composition, promote the formation of dynamic E-actin-based protrusions, and concentrate cell adhesion molecules to facilitate a complex morphogenetic event.

### Axons distinguish epithelial cell types

Although growing axons make direct contact with both periderm and basal keratinocyte membranes, we found that they dramatically reorganize the apical surface of basal keratinocytes, but affect the basal surface of periderm cells much less. These observations raise the question of how these cell-type-specific responses are achieved. One explanation is that axons can only affect membranes with particular lipid compositions. During differentiation, apical and basal epithelial membranes acquire distinct lipids, including the enrichment of PIP2 and Lo vs Ld membranes (Sampaio et al., 2011; Cao et al., 2012; Gerl et al., 2012), making this a plausible possibility. Basal and periderm cells may also differentially express distinct cell surface proteins mediating axon-skin cell recognition, including cell adhesion molecules, adhesion E-protein coupled receptors, or other receptors detecting secreted cues from axons. Finally, the two cell types may also possess distinct mechanical properties that impact cell-cell interactions, with apical membranes of basal cells being more deformable by axons than stiffer basal surfaces of periderm cells. These three possibilities are not mutually exclusive, and multiple factors may contribute to the basal cell-selective effects of axon contact.

Reciprocally, we found that basal cells remodel their membranes only in response to certain axons. Using live imaging, we found instances of axons inducing no microdomain formation or reorganization, even in the vicinity of axons that dramatically induced membrane remodeling. This selectivity presages the formation of ensheathment channels around some axonal segments, but not others (Yin et al., 2021). The cues mediating this specific axon recognition remain to be determined, but differential expression of cell surface cues on axonal membranes, or differences in mechanical properties of axonal segments, could play a role.

### Axons induce the reorganization of epithelial membranes

Previous work demonstrated that axon contact promotes the formation of microdomains on basal epithelial cells within minutes (Jiang et al., 2019). In this study, live time-lapse imaging revealed two mechanisms by which these microdomains form. In the first mechanism, pre-existing PIP2-enriched lipid microdomains coalesced to form larger domains beneath axons. In the absence of axons, the PIP2 reporter was present in patches, but these microdomains moved randomly within the membrane, indicating that axons induce their reorganization. In the second mechanism, PIP2 reporter-enriched microdomains sometimes formed anew, directly underneath axons. Both mechanisms imply that axons express cell surface molecules that rapidly induce remodeling of epithelial cell membranes through interactions with cell surface receptors. Those axonal proteins might be secreted or membrane anchored, but the fact that axon-induced basal cell microdomain remodeling was highly restricted to particular axon segments suggests that they likely act at short range. Furthermore, F-actin protrusions in these microdomains formed largely around axon shafts and not growth cones, whereas PIP2 lipid microdomains remodelled around both. These observations imply that the signals mediating neurite-induced epithelial remodeling act in a highly local manner, differing even along the length of a neurite. This localized mechanism could explain why ensheathment, which occurs over several days following axon growth into the skin, is neuron- and branch-specific.

In addition to molecular cues, local changes in cortical tension might also promote lipid microdomain association with axons. This idea was supported by an observation suggesting that lipid microdomains are sensitive to anisotropic changes in membrane tension: With live imaging studies, we captured basal cell division events that tugged on neighboring basal cells; membrane microdomains quickly disappeared when the non-dividing basal cell was stretched, but rapidly and specifically reformed next to axons after relaxation. This observation suggests that global changes in cortical tension can either inhibit (increased tension) or promote (relaxed tension) microdomain formation. This mechanism implies that particular axon segments could induce microdomain remodeling in underlying basal cells by locally relaxing cortical tension. A mechanical cue initiating microdomain reorganization around an axon would also ensure a localized response.

### Axon-associated microdomains have a distinct lipid composition

Membrane microdomains with a unique lipid composition had been previously proposed to be precursors of epidermal ensheathment channels, based on the observation that PIP2 reporters are enriched in both fish and fly neurite-associated domains (Jiang et al., 2019). However, without comparing this reporter to other membrane reporters, it was not possible to distinguish if these domains were biochemically distinct, or simply contained more membrane. We now provide quantitative evidence that axons rapidly promote the remodeling of epithelial membranes to create microdomains: A PIP2 reporter was significantly more enriched than a prenylated membrane reporter (mRuby-CAAX), implying that the accumulation of the PIP2 reporter in these domains is not explained simply by the presence of more membrane. The accumulation of PIP2 in these domains likely facilitates ensheathment, since inhibiting PIP2 production in Drosophila prevents neurite ensheathment by epidermal cells (Jiang et al., 2019).

In diverse contexts, membrane lipid composition at cell-cell interfaces, and at specialized membrane structures, exhibits a unique composition. For example, the formation of immune synapses, specialized junctions between antigen-presenting cells and antibody-producing B or T cells, requires the recruitment of signal transduction machinery into cholesterol- and sphingolipid-enriched Lo membranes (Dupre et al., 2002; Hiltbold et al., 2003; Nika et al., 2006; Zuidscherwoude et al., 2014). Clustering cell surface receptors within discrete membrane nanodomains can promote signaling activity (Miceli et al., 2001; Cambi et al., 2006; Jaumouille et al., 2014; Martinez-Munoz et al., 2018). To determine if similar Lo domains facilitate axon-epithelial cell communication, we measured the relative enrichment of membrane reporters that preferentially associate with Lo or Ld membranes. Indeed, at every developmental stage, Lo reporters were enriched over Ld reporters at axon contact sites, and this enrichment increased as axon-epithelial contacts matured.

PIP2- and Lo-enriched axon-associated domains form during initial axon contact, and a subset of these axon-associated microdomains give rise to ensheathment channels. We propose that Lo membranes may facilitate ensheathment channel formation in several ways. First, similar to immune synapses, they may concentrate signaling molecules that enable cell-type specific recognition between axons and epithelial cells. Second, a distinct lipid environment may promote membrane curvature around axons. Lipid rafts are associated with several highly curved membrane structures, including caveolae (Lamaze et al., 2017), viral budding sites (Sengupta et al., 2019), filopodia (Scorticati et al., 2011) and myelin sheaths (Yurlova et al., 2011). In the case of viral budding sites, acquisition of membrane curvature requires the partitioning of raft-enriched membrane (Sengupta et al., 2019). Finally, in addition to promoting cell-type recognition and membrane curvature, lipid rafts could recruit proteins that carry out later steps of ensheathment, including the recruitment of F-actin-binding and junction proteins. Thus, unique membrane microdomains have the potential to orchestrate several different activities that culminate in the formation of axon ensheathment channels.

### Axon-associated lipid microdomains are sites of F-actin protrusion

In previous work, we found that F-actin reporters accumulate at ensheathment channels in both flies and zebrafish (Jiang et al., 2019), suggesting that F-actin might facilitate cell shape changes associated with ensheathment and/or localize to autotypic adherens junctions. In this study, using live imaging, we found that F-actin associates dynamically with PIP2-enriched axon-associated microdomains at much earlier stages--soon after axons contact epithelial cells--and forms dynamic protrusions that selectively wrap around axons. Since several actin regulatory proteins, including Rho family GTPases, associate with PIP2 (Janmey et al., 2018), we propose that PIP2 enrichment at these microdomains promotes F-actin polymerization. Our observation that PIP2 microdomains can interact with both growth cones and axon shafts, whereas F-actin protrusions formed largely around axon shafts, suggests a stepwise maturation of axon-associated microdomains.

What is F-actin doing at these domains so early? Essential signaling complexes cluster in membrane microdomains in an actin-dependent manner(Dupre et al., 2002), and F-actin formation drives the lateral coalescence of these microdomains toward immune synapses (Dupre et al., 2002; Bolger-Munro et al., 2019). Thus, actin-enriched membrane microdomains may represent a generalized mechanism by which one cell recognizes another cell.

Microdomains at the basal cell surface could cluster proteins involved in axon-skin cell signaling and/or ensheathment channel formation, and F-actin dynamics could promote their coalescence underneath axons. F-actin protrusions may form as a result of epithelial membrane invagination around axons, similar to the actin-based pseudopods that form at nascent phagosomes. Later, during autotypic junction maturation, F-actin likely associates with adherens junctions. It will be interesting to determine the requirements of the actin cytoskeleton in the remodeling of membrane microdomains around axons, and identify the membrane proteins that accumulate in these microdomains.

### Axon-associated lipid microdomains enrich junction components to sequester them into ensheathment channels

We have identified epithelial cadherins as one class of membrane proteins enriched early in lipid microdomains. These microdomains progressively recruit cadherins as they mature, even in the absence of axons. Thus, axons do not themselves attract epithelial cadherins, but rather promote the remodeling of independently formed cadherin-enriched microdomains.

Actin-based protrusions containing cadherins initiate adherens junction formation between keratinocytes (called adhesion zippers; (Vasioukhin et al., 2000) and maintain stable adhesions by interdigitating membranes (called microspikes; (Li et al., 2020), suggesting that the protrusions we see at sites of axon contact may be related to these structures. During axon branching into the epidermis, lipid microdomains are dynamic, unstable structures, making transient contacts with periderm membranes and axons. Around axons, adhesion-zipper-like structures might promote adhesion of invaginating membranes as they wrap around axons, forming an autotypic adherens junction. By contrast, microdomains that do not associate with axons mature into stable periderm-basal cell adhesions. We found that lipid microdomains undergo a transition from dynamic to highly static membrane domains between 1 and 4 dpf.

This transition coincides with the gradual recruitment of cadherins. As cadherins are progressively enriched, it may be more likely that microdomains will be captured into stable adhesions through *trans* interactions.

The enrichment of PIP2 and raft-associated reporters at maturing ensheathment channels could be both a cause and a consequence of cadherin recruitment to axon-associated membrane domains. During epithelial and axo-glial junction formation, junction-associated proteins increasingly associate with Lo membrane (Honke et al., 2002; Schafer et al., 2004; Susuki et al., 2007; Resnik et al., 2011; Guillaume et al., 2013; Kurrle et al., 2013; Stahley et al., 2014; Lewis et al., 2019). For some membrane proteins, such as the desmosomal cadherin Desmoglein, partitioning into Lo membrane is essential for proper junction formation (Lewis et al., 2019).

Thus, we hypothesize that Lo membrane enrichment at axon-associated membrane domains is required for autoypic junction formation. At mature ensheathment channels, we found that acute depletion of membrane cholesterol reduced E-cadherin recruitment, suggesting that maintenance of E-cadherin around axons depends directly or indirectly on cholesterol-rich Lo membrane. Cadherin clustering in cholesterol- and sphingolipid-enriched membranes could provide a sorting cue at the trans Golgi network for cadherins (Cao et al., 2012), promoting their targeting to microdomains. The stronger enrichment in early microdomains of Desmocollin-2-like compared to E-Cadherin could reflect a stronger or more direct involvement of lipid sorting for the desmosomal cadherin relative to an adherens junction cadherin (Lewis et al., 2019).

Additionally, PIP2 recruits the Exocyst vesicle tethering complex (Liu et al., 2007); thus, microdomains may promote the docking of post-Golgi vesicles carrying Cadherins. Increased exocyst-dependent membrane delivery could also be important for increasing the surface area of membrane domains to wrap axons (Silva-Rodrigues et al., 2020). Local changes in PIP2 levels may generate second messengers that modulate intracellular calcium, which in turn regulates vesicle fusion. Finally, a highly ordered lipid environment could promote the lateral clustering of cadherins, increasing the avidity of trans cadherin interactions to promote junction formation (Zhang et al., 2009).

Most epithelial surfaces in our body are innervated, yet the overwhelming majority of studies on epithelial junction maturation involve systems devoid of sensory neurites. Thus, little is known about the effects of sensory axons on junction remodeling. Signals from parasympathetic and sympathetic neurons have been shown to regulate multiple steps of epithelial tubulogenesis, including branch formation (Bower et al., 2014; Nedvetsky et al., 2014) and apical membrane expansion (Nedvetsky et al., 2014). While previous work has focused on the role of neurotransmission and secreted peptides on epithelial morphogenesis--in some cases affecting epithelial properties through non-cell autonomous processes (Magnon et al., 2013)-- this work demonstrates how neurites can influence basic epithelial properties through direct physical interactions. We found that axon growth into the epidermis inhibits heterotypic junction formation by recruiting cadherin-enriched microdomains away from cadherin-rich periderm membranes and into autotypic junctions. Thus, by influencing the distribution of lipids, cytoskeletal elements, and cadherins, somatosensory axons exert an important influence on the distribution of cellular components whose polarized localization is critical for epithelial morphogenesis and homeostasis. Junctions are not merely spot-welds between cells but signaling centers; thus, the axon-mediated redistribution of junctions and cytoskeletal elements could have profound consequences on mechanical properties and mechanosensitive signaling in epithelia.

## Acknowledgments

We thank Kaitlin Ching for comments on the manuscript, and Son Giang and Linda Dong for excellent fish care. This work was funded by National Institutes of Health Dermatology T32 grant T32AR071307 to J.B.R. and National Institutes of Health grant R01AR064582 to A.S.

## Materials and Methods

### Zebrafish husbandry and microinjections

Zebrafish (Danio rerio) were raised at 28.5°C on a 14-h/10-h light/dark cycle. Embryos were raised at 28.5°C in embryo water composed of 0.3 g/L Instant Ocean salt (Spectrum Brands) and 0.1% methylene blue. Zebrafish lines used were Tg(*p63:*GAL4VP16; UAS: EGFP-PLCγ-PH) (Rasmussen et al, 2015); Tg(*krt5:GAL4*) (Rasmussen et al., 2015); Tg(*p63:*GAL4VP16; UAS: LifeAct-GFP) (Rasmussen et al., 2015); *cdh1*-TdTomato (Cronan et al., 2018); Tg(*krt5:*myrpalm-mCherry) (this work); Tg(*isl1:*GAL4VP16; UAS:EGFP) (Rasmussen et al., 2015); Tg(*isl1:*LexA; LexAop:TdTomato) (Rasmussen et al., 2015); Tg(*neural beta tubulin:DsRed*)(Peri and Nüsslein-Volhard, 2008).

Fertilized eggs were collected at the 1-cell stage for injections. ∼1 ng of the *ngn1* MO (5’ -ACGATCTCCATTGTTGATAACCTGG-3’; Andermann et al., 2002) or a standard negative control morpholino (5’-CTCTTACCTCAGTTACAATTTATA-3’; GeneTools) were injected into the yolk. Morpholinos were diluted in autoclaved double deionized water to working concentrations of 1 ng/nl. For plasmid DNA injections, ∼15-20 pg of DNA was co-injected with ∼25 pg *tol2* mRNA to facilitate recovery of somatic clones. Injection volumes were calibrated in drops of mineral oil on a slide micrometer.

### Molecular Biology

#### Generation of Lo and Ld membrane reporters

The myrpalm (mp) reporters were constructed according to sequences described in (Zacharias et al., 2002). Briefly, a forward primer containing the sequence 5’-C<u>ATG</u>GGATGTATTAATAGTAAGCGAAAGGAT-(gene specific sequence)-3’ was used to add the consensus sequence for the myristoylation and palmitoylation modifications at the N-terminus of the coding sequences for mEGFP, mApple, and mCherry, followed by TA cloning into pCRII-TOPO. cDNAs encoding the myrpalm-modified fluorescent proteins were subcloned into pDONR221-MCS-T2A for use in the Gateway cloning system (Kwan et al., 2007). See Supplemental Methods table for list of primers and plasmids used for cloning.

To make mp-mEGFP (A206K, L221K, L223K), site-directed mutagenesis was performed on pCRII-mp-EGFP using a QuikChange Lightning mutagenesis kit (Agilent) to introduce two further mutations, L221K and L223K, which eliminate any tendency of EGFP (A206K) to oligomerize. See Supplemental Methods table for list of primers and plasmids used for cloning.

To generate a GPI-anchored reporter, a codon-optimized signal sequence from human ER-resident p23 was inserted into the AgeI site of p-sfGFP-N1 (Addgene #54737). p23ss-sfGFP lacking the stop codon was PCR amplified and flanked with a 3’ NheI site and TA cloned into pCRII-TOPO vector. The introduced NheI site and a backbone EcoRV site in pCRII-TOPO was used to insert a codon-optimized C-terminal GPI signal from rat Neurotrimin (Uniprot Identifier NTRI_rat; amino acid sequence: N-AVSEVNNGTSRRAGCIWLLPLLVLHLLLKF*-C) (Galian et al., 2012). The resultant full-length p23ss-sfGFP-GPI was then inserted into pDONR221-MCS-T2A. See Supplemental Methods table for list of primers and plasmids used for cloning.

To label Ld membrane, a codon-optimized signal sequence with an N-terminal G_5_S linker sequence (encoding: N-DGKKKKKKSKTKCNLL*-C, derived from the Rap1b CAAX box) (Pyenta et al., 2001; Sengupta et al., 2019) was added to the C-terminus of mEGFP or mApple. mEGFP-gg and mApple-gg coding sequences were flanked with 5’ attB2r and 3’ attB3 recognition sequences to make p3E-mEGFP-gg and p3E-mApple-gg entry clones. See Supplemental Methods table for list of primers and plasmids used for cloning.

Sequence-validated pME-mp-mApple-T2A, pME-mp-mEGFP-T2A, and pME-p23ss-sfGFP-GPI clones were then used in LR Gateway recombination reactions with p5E-10xUAS or p5e-krtt1c19e (Rasmussen et al., 2015), and p3E-mEGFP-gg or p3E-mApple-gg, into the pDestTol2CG destination vector (Kwan et al., 2007). The p5E-krtt1c19e entry clone contains a promoter fragment from the *keratin type 1 c19e* gene that drives expression in basal keratinocytes (Rasmussen et al., 2015). A Kozak sequence (GCCGCCACCATGG) was included around the start codons for all expression constructs.

#### Generation of dual EGFP-PLCγ-PH membrane reporters

A HindIII-EGFP-PLCγ-PH-KpnI fragment was PCR amplified and subcloned into pDONR221-MCS-T2A. The resultant pME-EGFP-PLCγ-PH-T2A plasmid was used with p5E-10XUAS (Kwan et al., 2007), p3E-mRubyCAAX(Kwan et al., 2007) or p3E-mApple-gg in LR recombination reactions into the destination vector pDestTol2CG. See Supplemental Methods table for primers used.

Competent *E. coli* were grown in LB broth supplemented with antibiotics under standard conditions. Plasmid DNA was purified using Qiagen Mini Prep kits. For purification of Dsc2l-GFP Bacterial Artificial Chromosome, a NucleoSnap Plasmid Midi kit (Takara Bio) was used. PCR reactions were carried using Phusion DNA polymerase or Taq DNA Polymerase using commercial buffers (New England Biolabs). Reactions were run on a BioRad C1000 Touch thermal cycler. PCR products were either cleaned up using DNA Clean & Concentrator kits (Zymo Research) or were gel-purified using aGel Purification kit (Zymo Research). Plasmid DNA was transformed into OneShot Top10 competent cells (Invitrogen). Oligonucleotides were synthesized by Integrated DNA Technologies.

### Generation of Tg(*krt5*:mp-mCherry) transgenic fish

An attB1-mp-mCherry-attB2 PCR product was recombined into the pDONR221 vector. The resultant pME-mp-mCherry was used in a LR Gateway recombination reaction in combination with p5E-*krt5*, p3E-polyA entry clones and the pDestTol2CG destination vector. ∼15 pg of pDest-*krt5:*mp-mCherry-polyA was co-injected with ∼20 pg *tol2* mRNA into the ooplasm of WT AB oocytes at the 1 cell stage. F0 adults were crossed to WT fish to screen for founder fish that segregated mp-mCherry expression in the periderm of the F1s. 5 founders were identified to establish transgenic lines. For the experiments described in this paper, a single founder with ∼30% germline transmission was crossed to other transgenic lines for live imaging. See Supplemental Methods table for list of primers used.

### Microscopy and live imaging

For end-point imaging, live embryos or larvae were anesthetized in Tricaine and mounted in 1.2% low-melt agarose with their dorsal side against the coverslip to image the epidermis covering the head. Immobilized embryos were bathed in embryo water containing Tricaine within an imaging chamber formed by custom-made plastic rings sealed onto coverslips using vacuum grease (O’Brien et al., 2009). Imaging was performed using a Zeiss LSM 880 confocal microscope equipped with 488 nm (<0.5-1% laser power) and 561 nm excitation lasers (1-2% laser power). ∼10-25 um z-volumes were acquired using a 63x oil immersion objective and a 2-3x scan zoom.

For live imaging, manually dechorionated 1 dpf embryos were anesthetized in Tricaine and mounted as above. Imaging was performed using a Zeiss LSM 880 confocal microscope equipped with 488 nm (<0.5-1% laser power) and 561 nm excitation lasers (1-2% laser power). Slides were mounted on a stage heater heated to 28.5C. Z-stacks were acquired every 3-7 min for ∼5-7h. Movies were acquired using either a 20x air objective or 40x oil immersion objectives with 2-3x scan zoom.

### Cholesterol depletion with methyl-beta-cyclodextrin (MBCD)

Individual 4 dpf Tg(*p63:*GAL4VP16;UAS:EGFP-PLCγ-PH); *cdh1-*TdTomato embryos were placed in multiwell culture plates and incubated in 1 mL of vehicle (0.5% DMSO in isotonic Ringer’s), 5 mM MBCD (in 0.5% DMSO/Ringer’s diluent; Cayman Chemical), or 10 mM MBCD (in 0.5% DMSO/Ringers diluent) solution for 10 minutes. Embryos were then anesthetized using Tricaine in embryo water for ∼1 min. Embryos were rapidly mounted in 1.2% low melt agarose and imaged by confocal microscopy using 488 and 561 nm excitation wavelengths. Z-stacks for each embryo were collected. Methyl-beta-cyclodextrin solutions were prepared fresh the day of the experiment.

### Image Analysis

Image quantitation was done in ImageJ using maximum projected images. To quantify membrane reporter enrichment in microdomains and ensheathment channels, a ∼1.5-2 um linear ROI was drawn orthogonally across a microdomain or ensheathment channel to obtain intensity profiles. The ratio of the maximum Lo reporter intensity inside to the intensity outside was used to determine relative Lo enrichment. Ld reporter intensities at the same positions along the linear ROI were used to determine the relative Ld enrichment. >5 measurements were taken per basal cell clone. An identical approach was used to measure relative enrichment of Lo and Ld reporters in isolated microdomains. Relative enrichment values were averaged within individual clones to plot average relative intensities. To visualize cadherin recruitment to microdomains, intensity values along the linear ROI were normalized to the mean fluorescence intensity in a region encompassing both the microdomain and the surrounding non-microdomain membrane to produce plots of normalized intensities, allowing semi-quantitative comparison of cadherin recruitment across hundreds of microdomains with different transgene expression levels.

To quantify PIP2 lipid microdomain dynamics, ∼1 min interval movies were made in 1 or 4 dpf Tg(*p63:*GAL4VP16; UAS: EGFP-PLCγ-PH) zebrafish. To minimize image drift, movies were stabilized to the second frame with the ImageJ “Image Stabilizer” plugin. PIP2-rich microdomains in basal cells of 1 and 4 dpf control and *neurogenin1* morphants were selected as ROI with the polygon tool. Selected ROIs were duplicated to another window and cell borders were omitted from the analysis with the “Clear Outside” command. The brightness and contrast were automatically adjusted for the ROI and automatically thresholded with the “Default” algorithm. Frames within the movie were separated with the “Stack Splitter” command and overlap coefficients between the first frame and subsequent frames were extracted with the “Just Another Colocalization Plugin” (JACoP) in ImageJ.

### Transmission Electron Microscopy

Generation of transmission electron microscopy data was described in (O’Brien et al., 2012). The periderm-basal boundary was followed in TEM sections from ≥2 larvae per condition. Within a given image, the length of the periderm-basal boundary was traced and measured. The number of electron-dense junction plaques was counted and normalized to the length of the periderm-basal boundary. 19 sections were quantified for wild-type control larvae and 29 for *ngn1* morphants.

### Statistics

Statistical analysis was performed in Prism (v9.2.0) and R. Shapiro-Wilks tests were used to assess the normality of the distributions of data sets. One-way ANOVA or Kruskal-Wallis tests were used to compare the distributions across multiple timepoints, where appropriate. For pairwise comparisons of independent data, unpaired *t*-tests or Mann-Whitney U tests were performed. For paired data, paired *t-*tests or Wilcoxon signed rank tests were used. P values less than or equal to 0.05 were considered significantly different from the null hypothesis. See figure legends for specific statistical tests performed for each experiment.

## Movie Legends

**Movie 1. Axons induce remodeling of PIP2 membrane microdomains in basal cells** Time-lapse of 1 dpf Tg(*p63*:GAL4VP16; UAS:EGFP-PLCγ-PH); Tg(*neural beta tubulin:*DsRed) embryo. Axons are shown in magenta and basal cell PI(4,5)P2 in green. At the start of the movie, the apical surface of the basal cell contains isolated PIP2-containing membrane microdomains. Over time, PIP2 membrane microdomains form around axons and organize into axon-associated membrane domains strongly labeled by EGFP-PLCγ-PH. Cyan arrowheads show the growth cones of an axon making initial contact with the basal cell. Isolated PIP2 membrane microdomains appeared around the growth cone and coalesced underneath the axon. Red arrows point to an axon that had already grown over the basal cell at the start of the movie. Isolated microdomains formed along this axon before remodeling into larger axon-associated membrane domains. Images were acquired every 3.5 min.

**Movie 2. PIP2 membrane microdomains are dynamic and reversibly associate with axons** Time-lapse of 1 dpf Tg(*p63*:GAL4VP16; UAS:EGFP-PLCγ-PH); Tg(*neural beta tubulin:*DsRed) embryo. Axons are shown in magenta and basal cell PI(4,5)P2 in green. After the growth cone passed out of the field of view, PIP2 membrane microdomains formed, disappeared, then formed again underneath the axon shaft. Images were acquired every 5 min.

**Movie 3. Axons have little overt effect on periderm membranes** Time-lapse of periderm PIP2 membrane at 1 dpf labeled with UAS:EGFP-PLCγ-PH transgene injected into Tg(*krt5:*GAL4FF); Tg(*neural beta-tubulin:*DsRed) embryo. As an axon grew underneath the periderm cell (cyan arrowhead), PIP2 membranes did not re-organize above the axon. Faint maze-like membrane pattern is from microridges on the apical surface of the cell. Images were acquired every 3.5 min.

**Movie 4. Axons occasionally induce the formation of scattered PIP2 membrane microdomains in overlying periderm cells** Time-lapse of periderm PIP2 membrane at 1 dpf labeled with UAS:EGFP-PLCγ-PH transgene injected into Tg(*krt5:*GAL4FF); Tg(*neural beta-tubulin:*DsRed) embryo. In this example, faint, scattered PIP2 microdomains formed above an outgrowing axon, but they did not coalesce into larger axon-associated membrane domains during the movie. Cyan arrowhead marks the advancing growth cone. Images were acquired every 3.5 min.

**Movie 5. Axons induce the lateral coalescence of lipid microdomains in basal cells** Time-lapse of 1 dpf Tg(*p63*:GAL4VP16; UAS:EGFP-PLCγ-PH); Tg(*isl1*:LexA; lexAop:TdTomato). Axons are labeled in magenta and basal cell PIP2 in green. Prior to the appearance of the axon, the apical surface of the basal cell is labeled by isolated PIP2 lipid microdomains. After the growth cone passes over the basal cell, lipid microdomains order along filopodial tips before coalescing around the axon shaft proximal to the growth cone. Images were acquired every 5 min.

**Movie 6. Axons are required for the reorganization of PIP2 lipid microdomains** Time-lapse of cells in 1 dpf Tg(*p63*:GAL4VP16; UAS:EGFP-PLCγ-PH); Tg(*neural beta tubulin:*DsRed) injected embryos with *ngn1* MO. In the absence of axons, basal cells still formed lipid microdomains, but they did not show any directed reorganization, remaining scattered across the apical surface. Images were acquired every 5 min.

**Movie 7. Axons can induce the *de novo* formation of lipid microdomains on the apical membrane of basal cells** Time-lapse of 1 dpf Tg(*p63*:GAL4VP16; UAS:EGFP-PLCγ-PH); Tg(*neural beta tubulin:*DsRed) embryo. 10 minutes after the growth cone passed over the imaged basal cell, lipid microdomains appeared directly underneath the growth cone and axon shaft. New microdomains formed around the axon shaft (cyan arrows), and further remodeled by fusing together into a contiguous axon-associated membrane domain. Images were acquired every 5 min.

**Movie 8. Rapid changes in membrane tension remodel lipid microdomains in basal cells** Time-lapse of cells in 1 dpf Tg(*p63*:GAL4VP16; UAS:EGFP-PLCγ-PH); Tg(*neural beta tubulin:*DsRed) embryos. PIP2 lipid microdomains were imaged in a basal cell adjacent to a cell (asterisk). As the dividing cell rounded up, it pulled on the adjacent basal cell, causing a rapid disappearance of lipid microdomains. Following relaxation of the imaged cell, lipid microdomains rapidly reformed around axons. Images were acquired every 5 min.

**Movie 9. Association of lipid microdomains with axons is axon-selective** Time-lapse of the same 1 dpf Tg(*p63*:GAL4VP16; UAS:EGFP-PLCγ-PH); Tg(*neural beta tubulin:*DsRed) embryo as in Fig 3B. Axons are shown in magenta and basal cell PIP2 in green. Movie shows an example of an axon crawling over a basal cell with little or no remodeling of lipid microdomains on the underlying basal cell surface. Images were acquired every 5 min.

**Movie 10. PIP2 lipid microdomains are sites of F-actin accumulation** Time-lapse of 1 dpf Tg(*p63:*GAL4VP16; UAS:EGFP-PLCγ-PH) embryo injected with a 10XUAS:LifeAct-mRuby plasmid. LifeAct signal is shown in magenta and basal cell PIP2 in green. Movie shows an example of transient F-actin accumulation at PIP2 lipid microdomains in basal cells. Images were acquired every 6 min.

**Movie 11. Growth cones do not induce F-actin protrusions in basal cells** Time lapse of 1 dpf Tg(*p63*:GAL4VP16; UAS:LifeAct-GFP); Tg(*neural beta tubulin:*DsRed) embryo. LifeAct is in green and axons in magenta. An axon crawled over a basal cell but induced no protrusion near the growth cone (cyan arrowhead). Images were acquired every 3 min.

**Movie 12. Basal cell F-actin dynamically wraps axon shafts** Time-lapse of 1 dpf Tg(*p63*:GAL4VP16; UAS:LifeAct-GFP); Tg(*neural beta tubulin:*DsRed) embryo. LifeAct is shown in green and axons in magenta. Filamentous LifeAct signal dynamically surrounded axon shafts. Images were acquired every 3 min.

**Movie 13. PIP2 lipid microdomains are sites of F-actin protrusion formation** Time-lapse of a 1 dpf Tg(*p63*:GAL4VP16; UAS:EGFP-PLCγ-PH) embryo injected with a 10xUAS:LifeAct-mRuby plasmid. PIP2 is shown in green, F-actin in magenta. The imaged basal cell formed bright EGFP-PLCγ-PH- and LifeAct-mRuby-labeled foci (cyan arrowheads) that progressed into extended protrusions labeled strongly by EGFP-PLCγ-PH. The LifeAct signal was strongest at the base of the protrusion. Images were acquired every 5 min.

**Movie 14. Axon-selective formation of F-actin protrusions** Time-lapse of a 1 dpf Tg(*p63*:GAL4VP16; UAS:LifeAct-GFP); Tg(*neural beta tubulin:*DsRed) embryo. LifeAct is shown in green and axons in magenta. While some axons (cyan arrowhead) induced dynamic LifeAct-GFP-labeled membrane microdomains, a nearby axon (red arrowhead) had no effect on basal cell F-actin. Images acquired every 3 min.

**Movie 15. Quantitative analysis of lipid microdomains in the presence and absence of axons** Time lapse movies of 1 dpf and 4 dpf Tg(*p63*:GAL4VP16; UAS:EGFP-PLCγ-PH) embryos injected with a control (Ctrl MO) or *ngn1* morpholino (ngn1 MO). Representative movies used to quantify lipid microdomain dynamics, as described in Fig 7 and Supplemental Fig 4. Images were acquired every 1 min.

## Supplemental Figure Legends

**Supplemental Figure 1.**
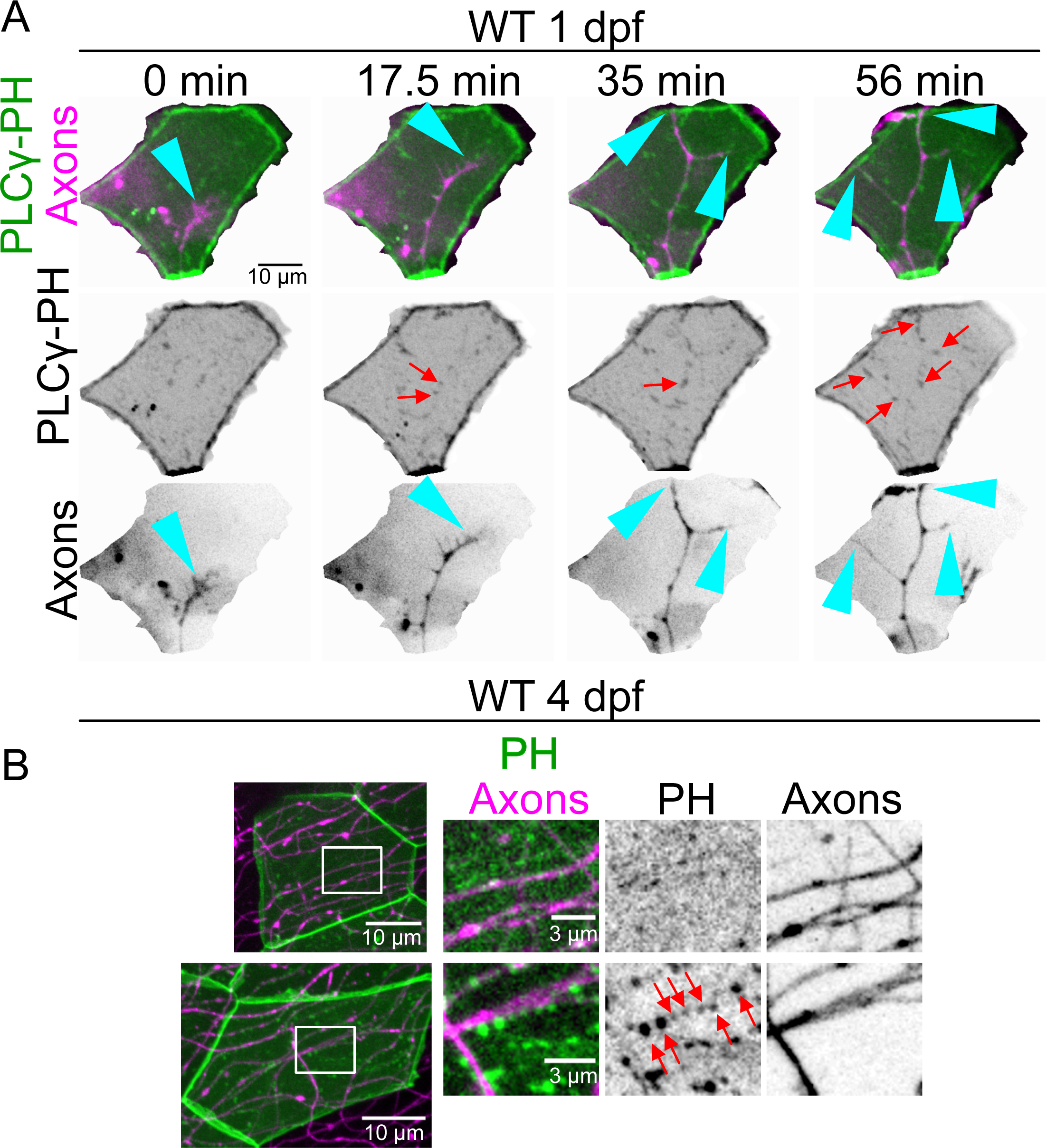
Periderm membranes do not markedly respond to axons during ensheathment stages. (A) Stills from a time-lapse movie of Tg(*krt5:*GAL4FF); Tg(*neural beta tubulin:*DsRed)1 dpf embryo injected with a UAS: EGFP-PLCγ-PH transgene to mosaically label PIP2 in periderm membranes. Scattered EGFP-PLCγ-PH-expressing membrane microdomains (red arrows) appeared above axons but did not coalesce into larger membrane domains. Cyan arrowhead marks growth cones crawling under the periderm cell. (B) By 4 dpf, when axons are ensheathed by PIP2-containing basal cell membranes, EGFP-PLCγ-PH either showed no distinct organization above axons (top cell) or occasionally formed discontinuous punctae above axons (bottom cell).

**Supplemental Figure 2.**
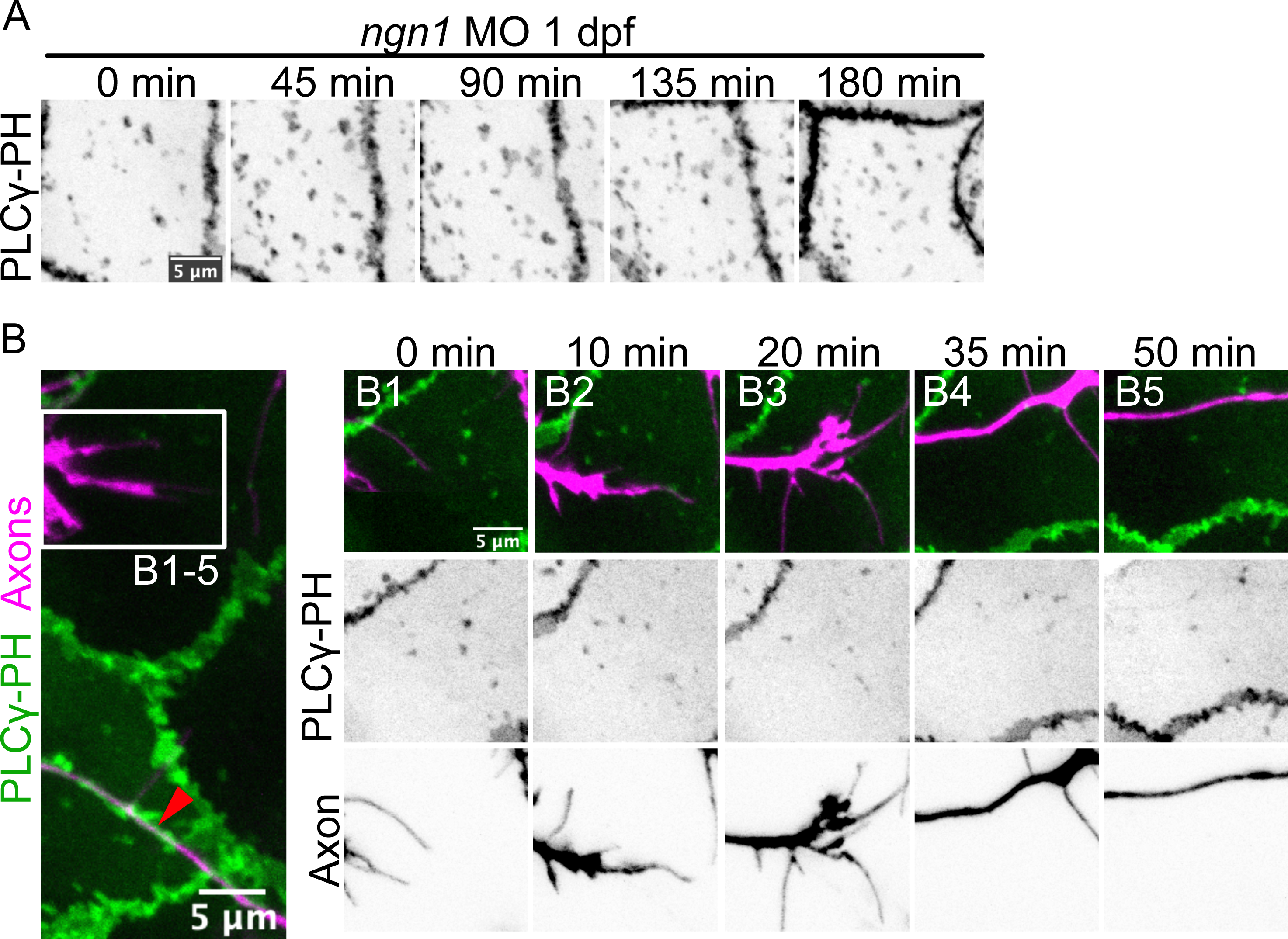
Epithelial membranes selectively respond to sensory axons. (A) Detail of the same region shown in Fig 3B from a Tg(*p63:*GAL4VP16;UAS:EGFP-PLCγ-PH; *neural beta tubulin:*DsRed) 1 dpf embryo. The axon marked by the red arrowhead rapidly induced *de novo* lipid microdomain accumulation (see Fig 3B1-5). However, a nearby axon branch (box) induced no lipid microdomain accumulation. (A1-5) Still from a time-lapse movie of boxed axon in A growing over a basal cell. (B) Stills from a Tg(*p63:*GAL4VP16; UAS:EGFP-PLCγ-PH; *neural beta tubulin:*DsRed) 1 dpf embryo injected with *ngn1* MO, resulting in a loss of sensory axons (not shown). Lipid microdomains formed normally but did not organize into coherent, linear domains.

**Supplemental Figure 3.**
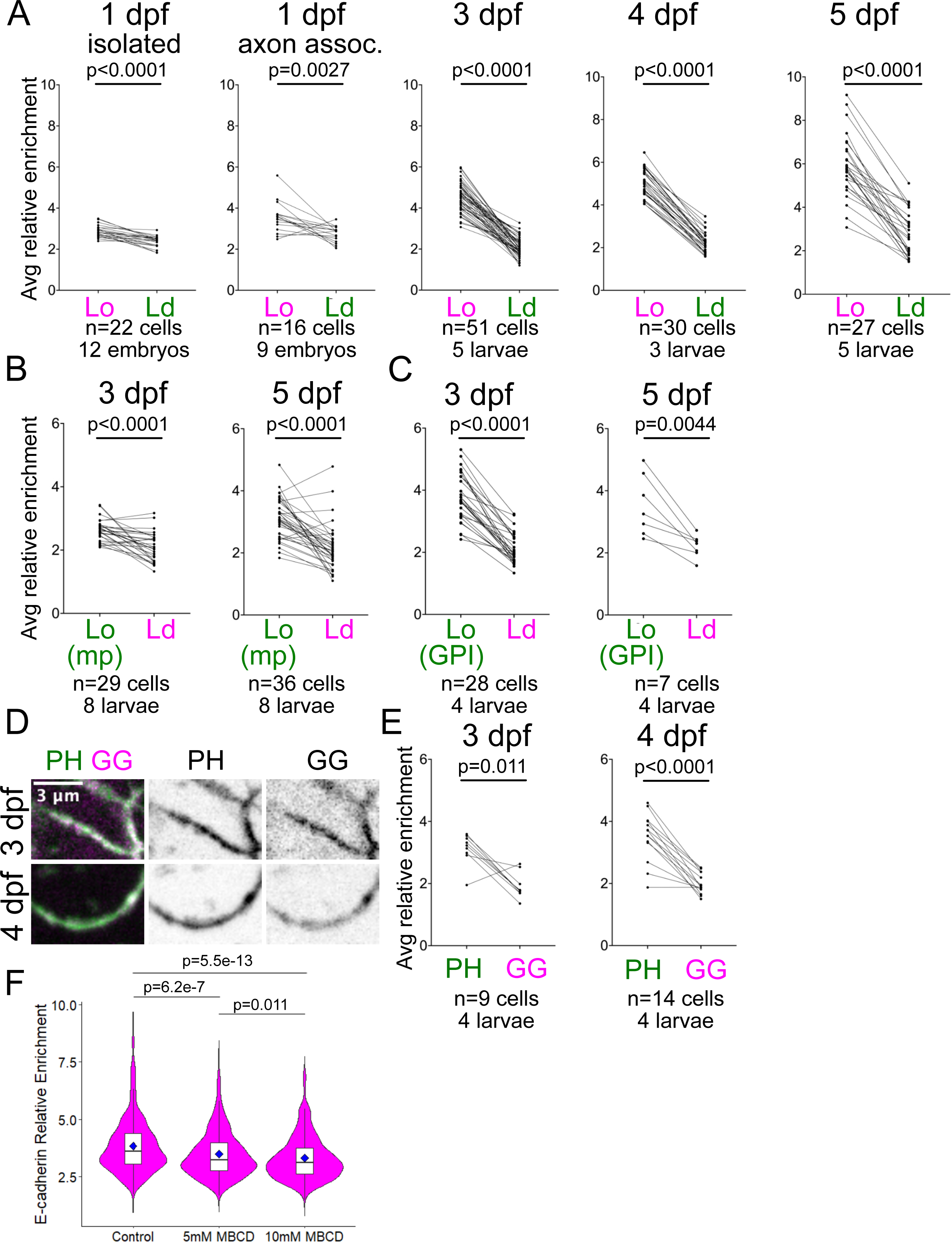
Axon-associated microdomains progressively enrich reporters for Lo, but not Ld, membrane. (A) Relative enrichment of mp-mApple and mEGFP-gg at axon-associated membrane domains from 1-5 dpf. Same data as in Fig 5D but represented as cell averages. (B) Relative enrichment of mp-mEGFP and mApple-gg at axon-associated membrane domains at 3 and 5 dpf. Same data as in Fig 5E but represented as cell averages. (C) Relative enrichment of sfGFP-GPI and mApple-gg at 3 and 5 dpf. Same data as in Fig 5E but represented as cell averages. (D) Representative image of axon-associated membrane domain in basal cell labeled with EGFP-PLCγ-PH-T2A-mApple-gg. (E) Relative enrichment of EGFP-PLCγ-PH (PH) and mApple-gg (GG) at 3 and 4 dpf. Data are shown as cell averages. (F) Relative enrichment of endogenous E-cadherin at axon-associated membrane domains from 3 dpf larvae treated with 0.5% DMSO in isotonic Ringer’s solution (control; n=87 cells, 16 larvae), 5mM methyl-beta-cyclodextrin in 0.5% DMSO (5 mM MBCD; n=94 cells, 16 larvae), or 10 mM MBCD in 0.5% DMSO (10 mM MBCD; n=93 cells, 15 larvae). Median is indicated by black line in the box plots. Blue diamond shows mean for each condition. P values computed using Mann-Whitney U tests.

**Supplemental Figure 4.**
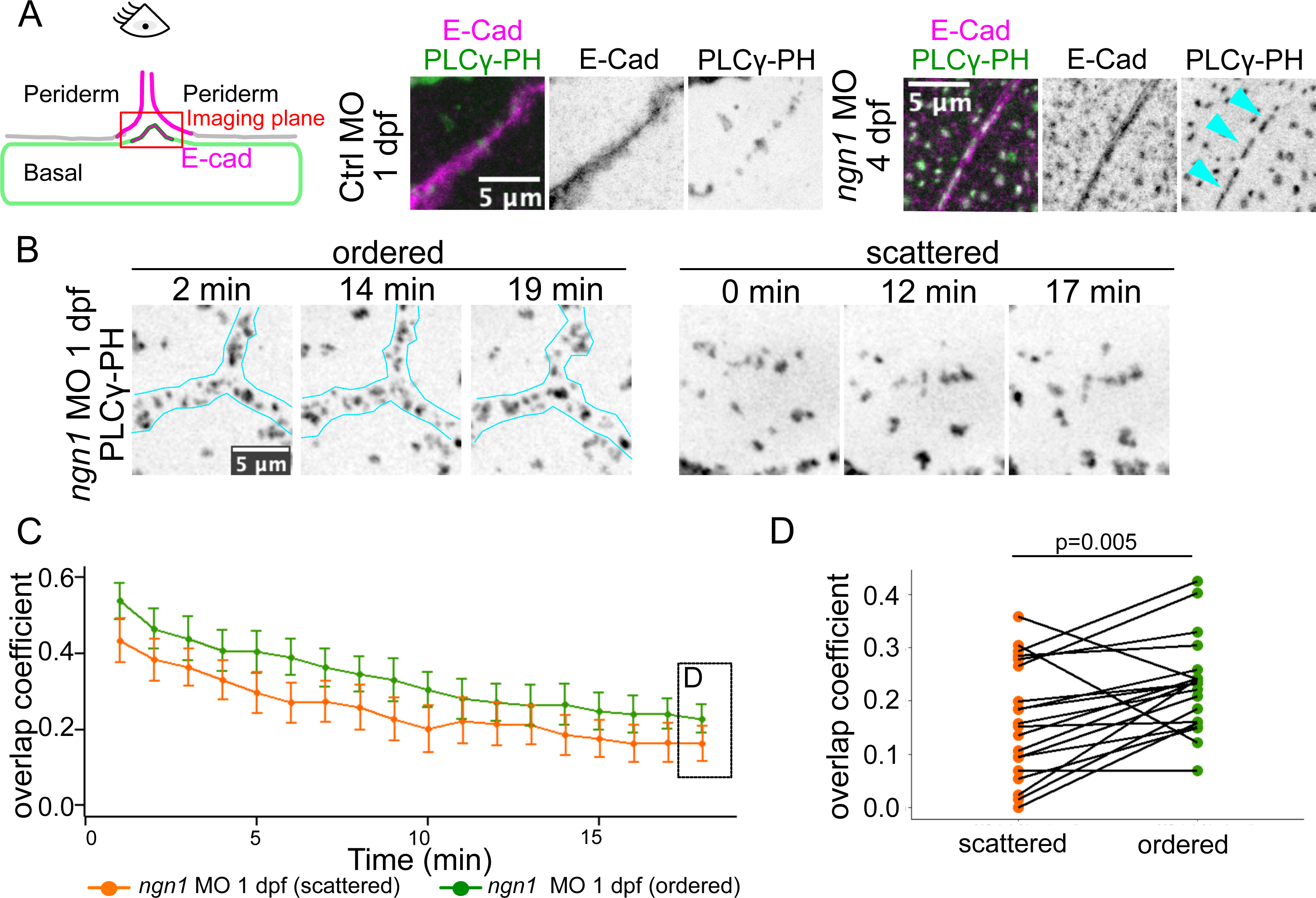
Interaction of lipid microdomains with cadherin-rich membranes suppresses their dynamics. (A) Cartoon schematizing the imaging of lipid microdomains in basal cells immediately underneath periderm-periderm borders. In wild-type control 1 dpf Tg(*p63:GAL4VP16:UAS:*EGFP-PLCγ-PH); *cdh1-TdTomato* embryos, PIP2 lipid microdomains (green) transiently ordered in register with cell-cell borders in the overlying periderm layer (labeled by endogenous E-cadherin reporter, magenta). In the absence of axons (*ngn1* MO), this transient ordering was more pronounced and resulted over time in the fusion of microdomains underneath cell borders (cyan arrowheads). (B) Representative stills from movies of ordered and scattered microdomains in 1 dpf *ngn1* morphants. Cyan lines demarcate the presumed position of periderm-periderm cell borders. (C, D) Comparison of dynamics of lipid microdomains that order underneath periderm cell borders (“ordered”) versus those unassociated with those regions (“scattered”) within the same basal cell. Ordering of lipid microdomains underneath E-cadherin-containing periderm membranes was associated with a suppression of their dynamics (p value computed using paired t-test; n=20 cells from 6 embryos).

**Supplemental Table 1.**
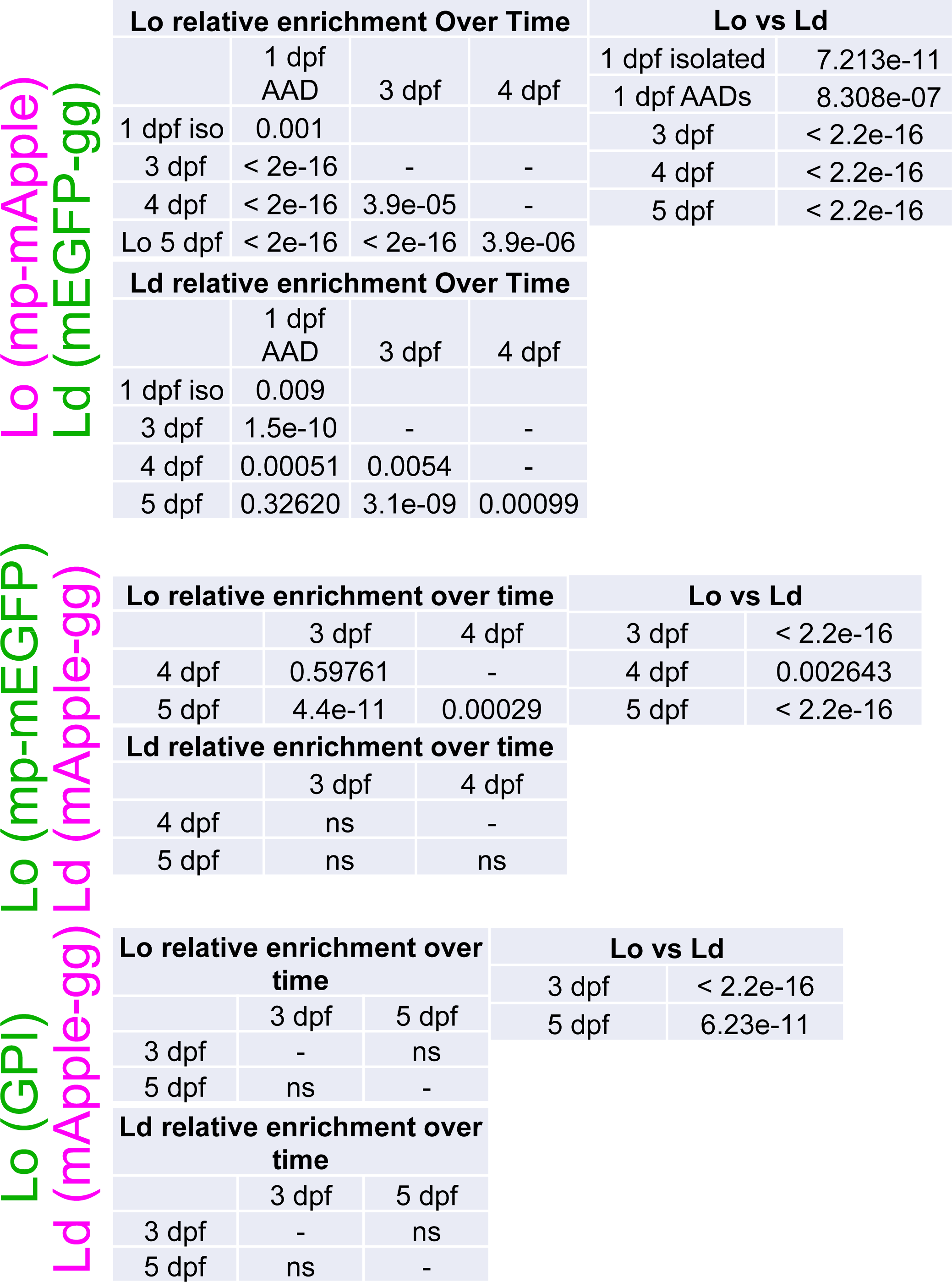
Summary of statistical analyses for Lo and Ld reporter enrichment. Table shows p values computed for Lo reporter enrichment over time and Lo versus Ld enrichment at each stage for all Lo and Ld reporters used in this study. See Fig 5 legend for details of statistical tests performed.

| Lo relative enrichment Over Time |  |  |  | Lo vs Ld |  |
| --- | --- | --- | --- | --- | --- |
|  | 1 dpf<br>AAD | 3 dpf | 4 dpf | 1 dpf isolated | 7.213e-11 |
| 1 dpf iso | 0.001 |  |  | 1 dpf AADs | 8.308e-07 |
| 3 dpf | < 2e-16 | - | - | 3 dpf | < 2.2e-16 |
| 4 dpf | < 2e-16 | 3.9e-05 | - | 4 dpf | < 2.2e-16 |
| Lo 5 dpf | < 2e-16 | < 2e-16 | 3.9e-06 | 5 dpf | < 2.2e-16 |
| Ld relative enrichment Over Time |  |  |  |  |  |
|  | 1 dpf<br>AAD | 3 dpf | 4 dpf |  |  |
| 1 dpf iso | 0.009 |  |  |  |  |
| 3 dpf | 1.5e-10 | - | - |  |  |
| 4 dpf | 0.00051 | 0.0054 | - |  |  |
| 5 dpf | 0.32620 | 3.1e-09 | 0.00099 |  |  |

Lo (mp-mEGFP) Ld (mApple-gg)
| Lo relative enrichment over time |  |  | Lo vs Ld |  |
| --- | --- | --- | --- | --- |
|  | 3 dpf | 4 dpf | 3 dpf | < 2.2e-16 |
| 4 dpf | 0.59761 | - | 4 dpf | 0.002643 |
| 5 dpf | 4.4e-11 | 0.00029 | 5 dpf | < 2.2e-16 |
| Ld relative enrichment over time |  |  |  |  |
|  | 3 dpf | 4 dpf |  |  |
| 4 dpf | ns | - |  |  |
| 5 dpf | ns | ns |  |  |

Lo (GPI) Ld (mApple-gg)
| Lo relative enrichment over time |  |  | Lo vs Ld |  |
| --- | --- | --- | --- | --- |
|  | 3 dpf | 5 dpf | 3 dpf | < 2.2e-16 |
| 3 dpf | - | ns | 5 dpf | 6.23e-11 |
| 5 dpf | ns | - |  |  |
| Ld relative enrichment over time |  |  |  |  |
|  | 3 dpf | 5 dpf |  |  |
| 3 dpf | - | ns |  |  |
| 5 dpf | ns | - |  |  |

